# Inferring gene expression models from snapshot RNA data

**DOI:** 10.1101/2022.05.28.493734

**Authors:** Camille Moyer, Zeliha Kilic, Max Schweiger, Douglas Shepherd, Steve Pressé

**Author notes:** These authors contributed equally (Camille Moyer, Zeliha Kilic, Max Schweiger).

## Abstract

Gene networks, key toward understanding a cell’s regulatory response, underlie experimental observations of single cell transcriptional dynamics. While information on the gene network is encoded in RNA expression data, existing computational frameworks cannot currently infer gene networks from such data. Rather, gene networks—composed of gene states, their connectivities, and associated parameters—are currently deduced by pre-specifying gene state numbers and connectivity prior to learning associated rate parameters. As such, the correctness of gene networks cannot be independently assessed which can lead to strong biases. By contrast, here we propose a method to learn full distributions over gene states, state connectivities, and associated rate parameters, simultaneously and self-consistently from single molecule level RNA counts. Notably, our method propagates noise originating from fluctuating RNA counts over networks warranted by the data by treating networks themselves as random variables. We achieve this by operating within a Bayesian nonparametric paradigm. We demonstrate our method on the *lacZ* pathway in *Escherichia coli* cells, the *STL1* pathway in *Saccharomyces cerevisiae* yeast cells, and verify its robustness on synthetic data.

## 2 Introduction

Quantitative measurements of RNA dynamics in populations of fixed cells and individual living cells have consistently revealed complex distributions of RNA counts and behavior across space, time, and individual cells, even for clonal cell populations [1]. Broadly, the term “gene expression variability” is invoked to explain these ubiquitous, variable, and complex RNA expression distributions. One of single-cell biology’s driving goals is understanding the molecular origin and downstream consequences of gene expression variability. For example, recent work has demonstrated that rare cells, identifiable only by transient fluctuations in RNA content compared to clonal sister cells, can drive drug-resistant cancer or maintain progenitor cells driving development [2–4]. While experimental methods can identify rare cells, robustly determining gene networks from discrete RNA counts across cells remains an open problem.

Fig. 1 shows examples of gene networks defined by their number of gene states, state connectivities, and associated rate parameters.

**Fig. 1.**
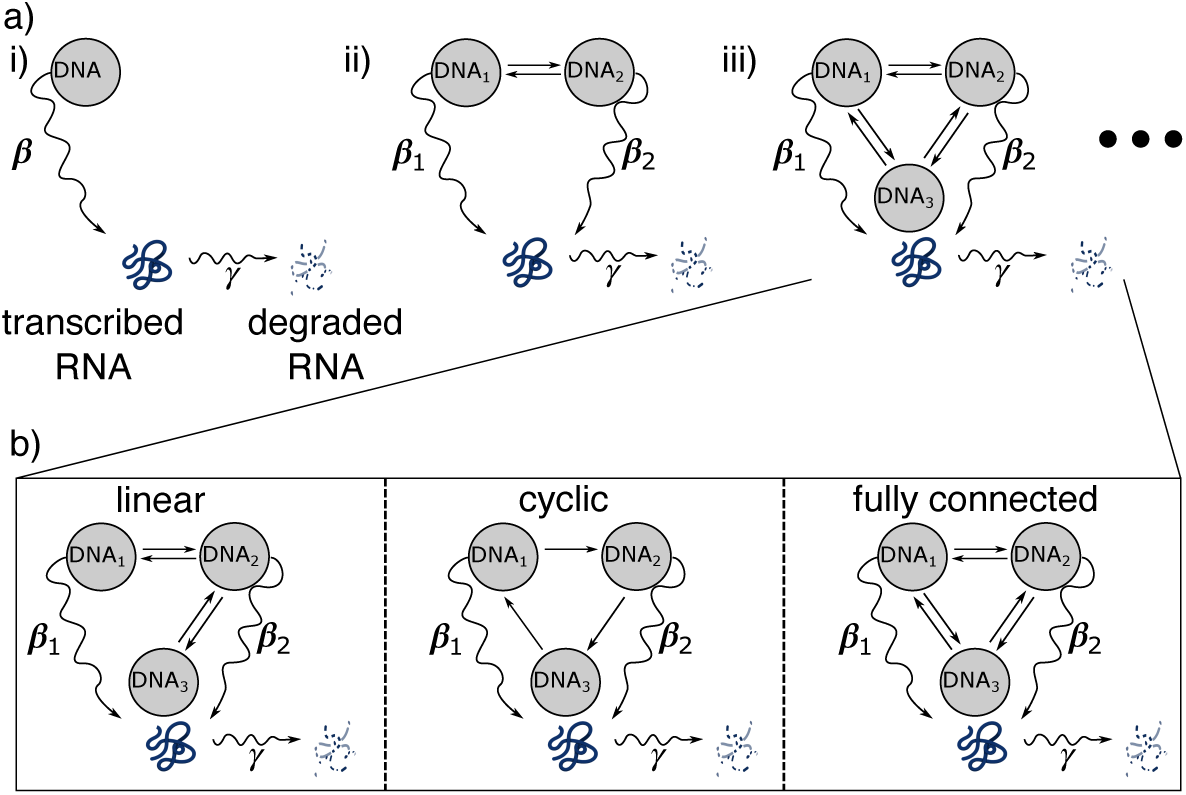
Cartoon of gene state models. Here we show a cartoon representation of one, two, and three state gene models (panel a). Each grey circle depicts an RNA production state that a gene may occupy, differentiated by its unique production rate. Straight arrows reflect possible transitions between gene states, and curved arrows depict RNA transcription (with rate β) or degradation (with rate γ). Panel b depicts models with a variety of transitions omitted. As we will see shortly, we will propose a method to infer gene networks (which includes gene state numbers and associated rates). As such, we will learn the ‘connectivity’ directly directly from the data.

Of the existing experimental methods providing discrete RNA counts, we focus on single-molecule RNA fluorescence *in situ* hybridization (smFISH [5, 6]). In particular, smFISH provides *snapshot* data consisting of independent fluorescent imaging assays performed on fixed samples at discrete time points, often following external stimuli. These assays yield the number and location of individual transcripts for a limited number of RNA species for individual cells and, as a consequence, direct insight into the molecular state of the cells or tissue at the time of fixation.

To help highlight the major challenges facing computational inference of gene networks from snapshot smFISH data, we consider the simplest gene network, shown in Fig. 1 a i), which consists of a single gene state. The one-state model predicts a Poissonian number of RNAs transcribed per time interval [7, 8]. A Poisson statistical expectation often disagrees with observation [9–16], and, as such, the perennial two-state model (Fig. 1 a ii) is conjectured [1, 10, 11, 13–21]. The two-state, or telegraph, model allows the gene regulatory network to transition between an inactive and active state. Despite the surprising range of behavior produced by the two-state model [21], it often fails to describe observed distributions in smFISH snapshot data. This, in turn, suggests the existence of gene states with intermediate RNA production rates. Immediately, challenges emerge on account of the many ways of connecting *N* ≥ 3 gene states (see Fig. 1 b for some examples). The additional reaction pathways may provide a better fit to existing data at the cost of predictive power. In fact model inference, which rigorously balances data description with predictive power, has yet to be achieved for this problem. In a number of previous attempts, metrics, including Poisson indicators [7, 9–11, 16, 22–25], crossvalidation [26], nonparametric regression [27–33], or information metrics which compare a truncated set of possible models [22, 23, 34, 35] are used to justify the introduction and network structure of additional gene states (Fig. 1 a iii). However, all such methods perform model selection and parameter inference separately and cannot therefore propagate error from inherently stochastic RNA counts into uncertainty over gene networks. As such, they fail to balance descriptive ability and predictive power in a statistically rigorous manner, and the relative probability over each proposed network, including gene states and associated parameters and thus connectivity of the gene states, given the data remains unknown.

In a further simplification, some of these methods altogether ignore the intrinsic stochasticity of RNA counts in favor of mass action formulations, which predict the temporal evolution of the mean number of RNA molecules per cell [7, 10, 11, 16, 24, 36, 37]. Mass action formulations are fundamentally insufficient for smFISH data, as RNA copies may be present at low numbers, rendering information on their copy number fluctuations vital toward extracting kinetic parameters [21]. Even methods which resolve this issue using the forward Kolmogorov equation (chemical master equation or CME), which gives the probabilistic temporal evolution of single-cell RNA counts [8, 35, 37–43], still lack robust methods to learn gene states.

Here, we propose a method to simultaneously arrive at a probability over gene states and their associated parameters, and thus connectivity, given the discrete RNA counts in smFISH snapshot data sets by meaningfully propagating inherent noise into the model estimation. We achieve this within the Bayesian paradigm, which allows us to draw samples from posterior probability distributions over gene networks. In order to construct these posteriors, we require priors over gene state numbers, necessitating the use of Bayesian Nonparametrics (BNPs) [44–48] and a likelihood over the data given model parameters, which we calculate using the CME. With a likelihood and nonparametric priors at hand, we simultaneously and self-consistently estimate model structures (number of gene states) alongside associated rate parameters and thus connectivities between states, as warranted by observed RNA counts per cell.

We demonstrate our method’s applicability by first testing it on synthetic data followed by two very different experimental systems: the *lacZ* pathway in *Escherichia coli* (*E. coli*) cells [49] and the *STL1* pathway in *Saccharomyces cerevisiae* (*S. cerevisiae*) yeast cells [50].

## 3 Results

We assume the availability of snapshot smFISH data containing RNA counts per cell, 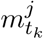, collected at time points, *t*_1:*K*_, and cells indexed as *j* = 1, …, *J*_*k*_. For simplicity, we denote the RNA counts from all cells at all time points 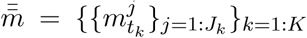. Using this information, our goal is to predict the transcriptional gene output, i.e., we infer both the gene states (that is, perform model selection) as well as infer the associated rate parameters and thus gene state connectivity (parameter inference) [10, 11, 26, 38, 39, 49–51].

Here we show our method’s ability to perform model selection and parameter inference for gene networks using both experimental and synthetic data. In BNPs, the model structure (in this case the number of gene states) is treated as a parameter, and thus inferred alongside all other parameters. The remaining parameters of interest are: the production rates, *β*_*l*_, for each gene state; the transition rates between various gene states, 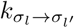 for *l* ≠ *l*^*′*^; and the RNA copy rate of degradation, *γ*. Since we are working within the BNP paradigm, our parameter estimates are drawn from fully joint posterior probability distributions over rates as well as gene states, learned simultaneously and self-consistently. Samples from these posteriors are displayed below in the form of histograms. Naturally, these histograms provide a complete assessment of each quantity’s value alongside their respective uncertainties.

First, we demonstrate the method’s robustness on synthetic data which replicates dynamics similar to experimental data sets, but covers a broader range of scenarios than are available experimentally. Subsequently, we show results for experiments on the *lacZ* pathway in *E. coli* cells [49] then the *STL1* pathway in *S. cerevisiae* yeast cells [50].

### 3.1 Robustness Analysis

#### 3.1.1 Number of states

In line with the current literature on gene expression [10, 39, 55–57], we tested our method on three different models consisting of one, two, and three gene states. In the two and three state models, one state has a production rate of zero.

Fig. 2 shows the results of the method for one, two and three gene state models. As we are working with synthetic data and ground truth is known, we can ascertain that the method is successful in placing substantial posterior probability on parameters close to ground truth.

**Fig. 2.**
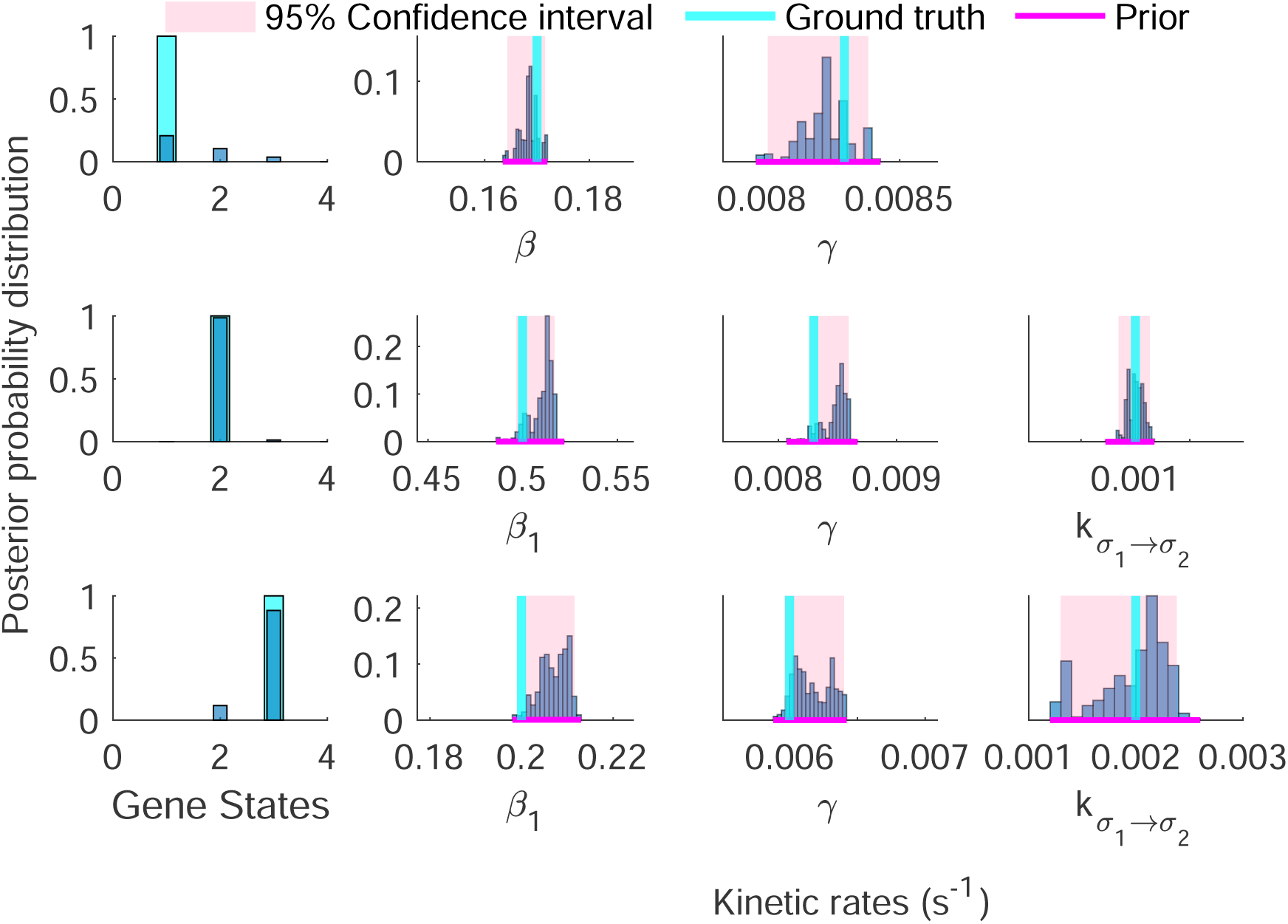
Accurate inference for a variety of gene networks from synthetic data. Here we show posterior distributions over: gene states (first column), production rates β_l_ (second column), degradation rates γ (third column), and transition rates 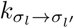 (fourth column). In the first row, we show distributions for a one gene state model, i.e., production at a single rate β and degradation at rate γ, and no transition rates. As desired, our posterior maximum closely matches the ground truth. We find similar results for the second and third rows, which depict two and three gene state models. Each synthetic data set is composed of 2000 cells observed per time point with 20 collection times evenly spaced over 1 hour [50]. Each data point was generated using the Gillespie stochastic simulation algorithm [52, 53]. Histograms of rates not shown here are included in Fig. S 12. For this and subsequent figures, we utilized a publicly available Matlab implementation of the Munkres assignment agorithm [54]. This code was used as a post-processing step, solely for the purpose of assigning labels to gene-states appropriately across MCMC iterations and only affects our figures.

Perhaps counter-intuitively for the simplest of gene networks, which should be easiest to infer, the posteriors over gene states are broader. The reason is subtle: models that estimate a greater number of gene states can approximate a one state model (but not vice versa) by having nearly identical production rates for each of the gene states. However, since the production rates of all possible gene states are not perfectly identical, our algorithm still favors the true number of states.

#### 3.1.2 Quantity of data

In Fig. 3, we test the method’s robustness with respect to data set size. For networks with two gene states, the method makes accurate inference across three orders of magnitude in the number of cells extracted per time point Fig. 3. While we find that the precision of estimates (breadth of posteriors) made by the model does scale with the quantity of data, the accuracy does not. In fact, the method makes accurate inference on the two gene state model provided data on only a handful of cells, with posterior probabilities whose breadth reflect the small quantity of data.

**Fig. 3.**
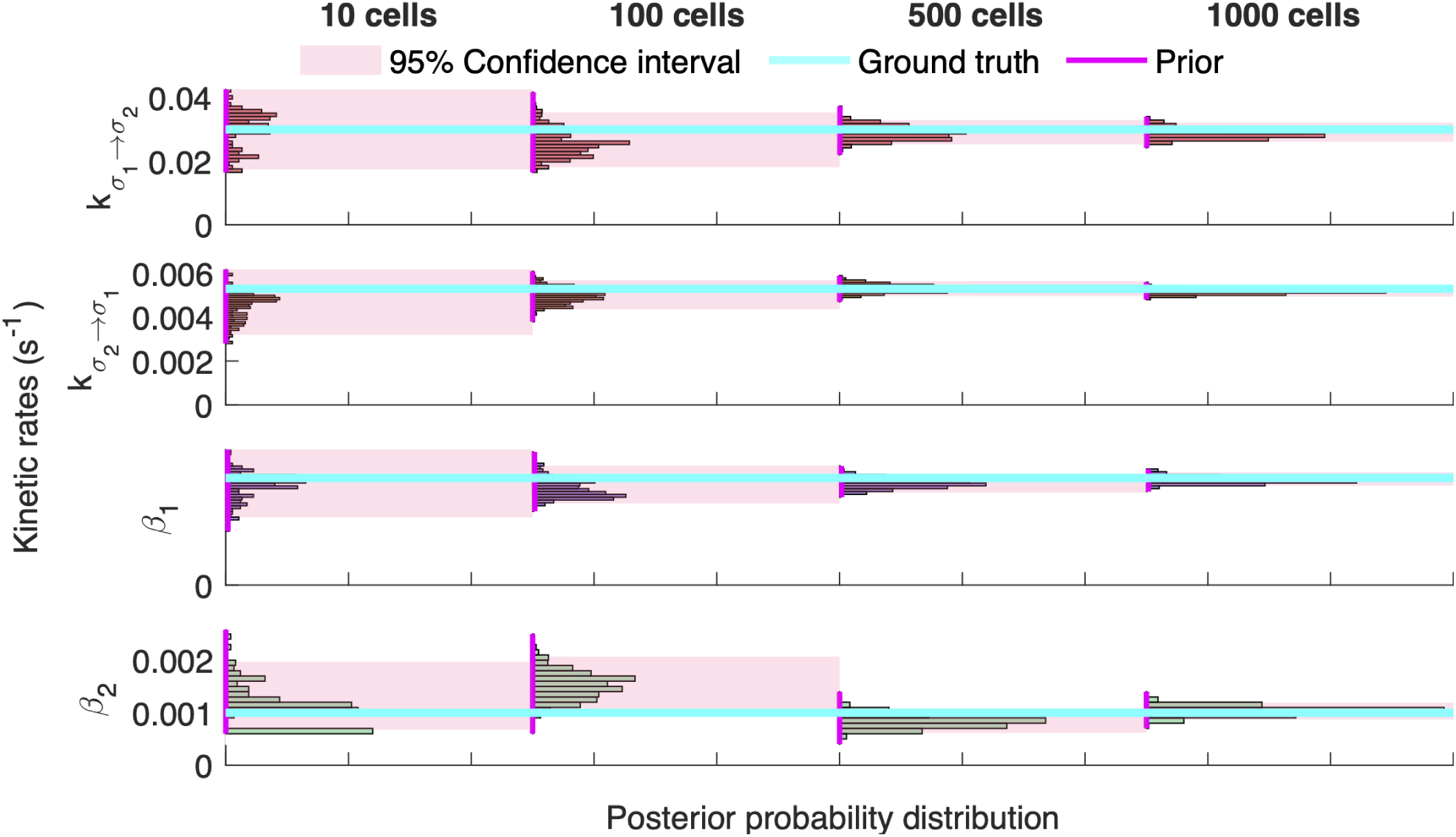
Sensitivity analysis: quantity of data. Here we show posterior distributions over: production rates β_l_, and transition rates 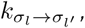, for networks with two gene states. Each row shows a different kinetic rate’s posterior becoming increasingly narrow as the quantity of data used in the analysis grows. As such, we find that our posterior maximum closely matches the ground truth, and confidence intervals, become markedly narrower as the quantity of data increases. Synthetic data sets are composed of 10, 50, 500, or 1000 cells observed per time point with 20 collection times evenly spaced over 1 hour, based on typical experimental procedures [50]. As before, each data point was generated using the Gillespie stochastic simulation algorithm [52]. Posteriors over the number of gene states are omitted for this and subsequent figures, as they place complete confidence on the correct number of gene states even with as few as 10 cells per time point.

For a detailed overview of the remaining robustness analysis of our method, we refer the reader to Section S 1.7.

#### 3.1.3 Synthetic Data Generation

Synthetic data used in Section 3.1 was generated using computer simulation based on Gillespie’s Stochastic Simulation Algorithm [52]. Details of the model are outlined in Section S 1.1, with inference of all parameters detailed in Section S 2.1.

### 3.2 Experimental Results

We analyzed smFISH RNA count data from *E. coli* [49] and *S. cerevisiae* [50] cells. For simplicity here, we calibrate the RNA molecules’ degradation rate using previously established results [39, 49] and demonstrate in Fig. S 2 that calibrating one rate, as expected, has the net effect of reducing uncertainty in the other rates inferred.

For greater detail regarding experimental conditions, we refer the reader to [49] (*E. coli*) and [50] (*S. cerevisiae*).

#### 3.2.1 E. coli

Here we demonstrate our simultaneous gene state number and parameter inference for the *lacZ* pathway in *E. coli* cells, grown in slow-growth media. Our results are shown in Fig. 4. In this figure, we also show the point estimates obtained in [49] (referred to as Wang et. al.) from maximum likelihood. To be clear, [49] posit a model (i.e., pre-specify gene states) and, given the model, learn parameters from the data.

**Fig. 4.**
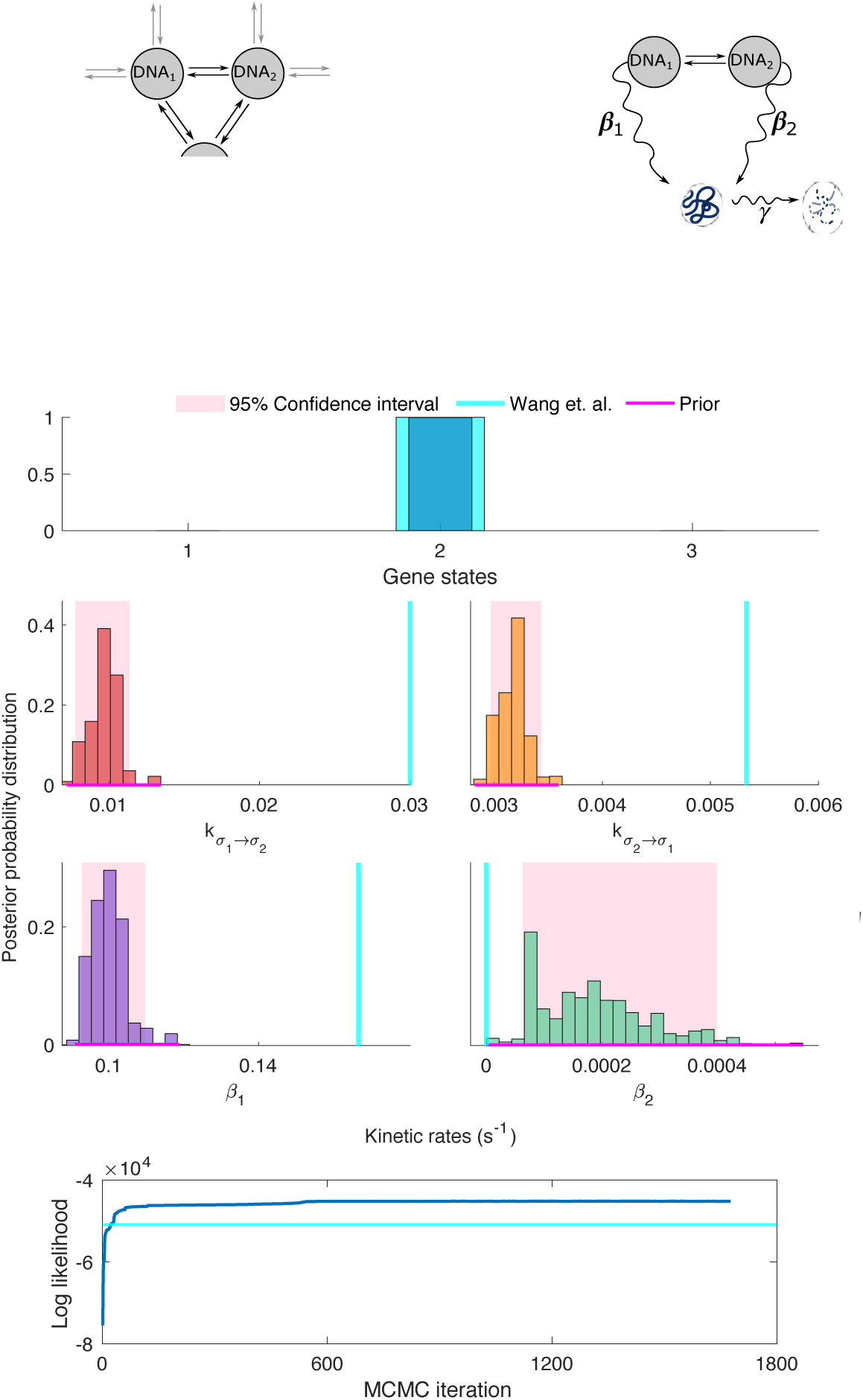
Inference on slow-growth E. coli data. Here we show results of analysis of the lacZ pathway in E. coli grown in glycerol at 30°C. Each panel represents the assigned posterior probability distribution for different model parameters, as compared to estimates from Wang et. al. [49], shown as vertical cyan lines. Pink shaded regions depict intervals in which 95% of MCMC samples lie. We recover a two gene state model with parameters which differ from those in [49] as outlined in Section 3.2.1. The bottom panel shows a trace of our method’s log likelihood surpassing that of [49] (depicted as a horizontal cyan line). See Fig. S 9 for joint histograms of rates shown here.

We find that the states they posit agree with what is learned directly from the data. In terms of parameters, we also find general agreement in the lowest production rate. However, we find disagreement of 70%, 40%, and 43% in parameters 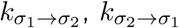 and *β*_1_ respectively, when comparing our maximum *a posteriori* (MAP) estimate to those reported in [49]. This disagreement highlights a core issue: even when learning only rates (and positing states by hand) [49] cannot efficiently sample their high dimensional posterior. To wit, we have employed Hamiltonian Monte Carlo (HMC) and Parallel Tempering (PT) for this reason, allowing our method to avoid becoming trapped in local maxima. As a result, we find that a comparison of likelihoods dramatically favors the parameters to which we have converged (by contrast to those reported in [49]). Indeed, we find that our MAP estimate (*θ*^*′*^) is more probable than that of [49] (*θ*) by a factor of 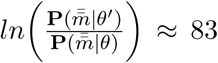. A trace of our method’s log likelihood surpassing that of [49] can be seen in Fig. 4.

Interestingly, despite the significant difference between *θ* and *θ*^*′*^, RNA count histograms across time points appear qualitatively similar; see Fig. S 3. This is expected as static histograms do not contain temporal information that we leverage in analyzing time-ordered snapshot data. By the same token, it highlights the limitations of assessing kinetic rates and gene states by comparing RNA histograms across time points to the method proposed herein.

Fig. 5 compares our learned model to that estimated in [49] for *E. coli* grown in a fast-growth medium (glucose at 37^*°*^*C*). Our full nonparametric method confidently infers three gene states, by contrast to [49], who assume two states. As we predict different models, a direct likelihood comparison is more difficult here than in the previous case.

**Fig. 5.**
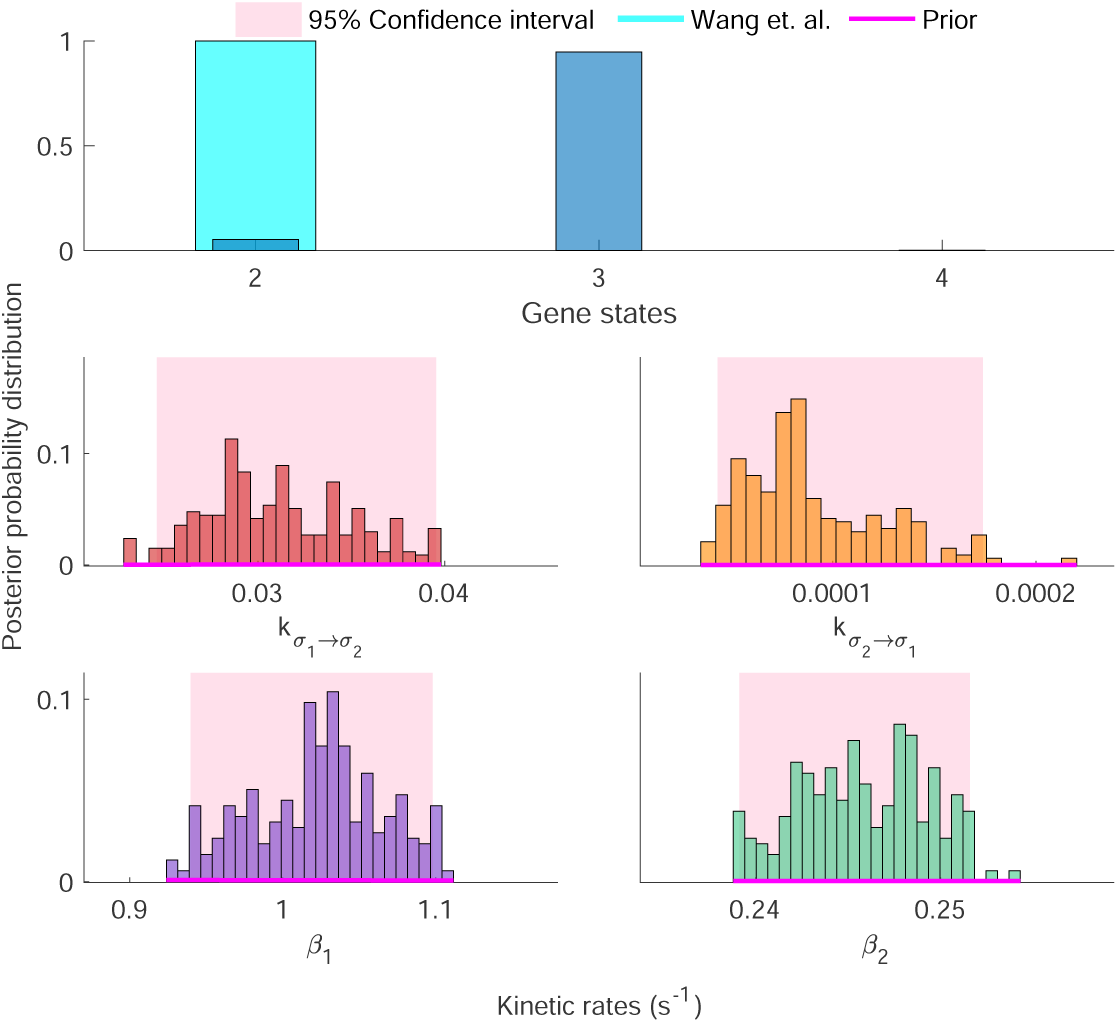
Nonparametric inference on fast-growth E. coli data. Here we show a subset of inferred rates for the lacZ pathway in E. coli grown in Glucose at 37°C. Here our method strongly favors three gene states. As with subsequent results for three gene state dynamics, we show rates of production β_1_, β_2_, and transition 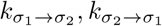 associated to the states with the two greatest production rates. Except for the number of gene states, the parameters inferred by Wang et. al. are omitted, as they result from an assumed number of gene states which differs from the number we estimate. Fig. S 5 further illustrates the point that histogram fitting may be an inadequate means for estimating rates. See Fig. S 10 for joint histograms of rates shown here.

However, in order to directly compare likelihoods, we restrict our method to infer parameters by imposing ourselves by hand a two state gene expression model, with one production rate fixed at zero (Fig. S 7). We find disagreement of similar order to that shown before: 78%, 67%, and 18% difference in parameters 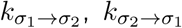 and *β*_1_ respectively. A ratio of likelihoods again favors our estimate 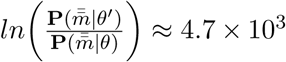. This again suggests the presence of local maxima that may lead to incorrect parameter value estimates even when assuming a simpler model with fewer states. Once more, this result underscores the need for simultaneous optimization methods such as our own.

#### 3.2.2 S. cerevisiae

Fig. 6 shows the results of model inference on the *STL1* pathway in *S. cerevisiae* cells. We compare our findings on gene states and parameters inferred to estimates of gene parameters alone (and gene states posited) from [39]. Our analysis confirms the four states of chromatin reorganization of the *STL1* gene in *S. cerevisiae* conjectured by previous methods prior to parameter estimation [26, 39]. However, as outlined in Section 2 and Section 3.2.1, this prespecification of gene state numbers may lead to parameter estimates which only locally maximize the likelihood for a given set of observations. Owing to our improved exploration of the space of models, we learn parameters (*θ*^*′*^, shown in Fig. 6) which we calculate to be more likely than those found in [39] by a factor of ln 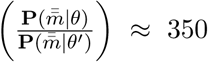 for *STL1* transcription. See Fig. S 8 for a comparison of predictive distributions of the type referred to in Section 3.2.1.

**Fig. 6.**
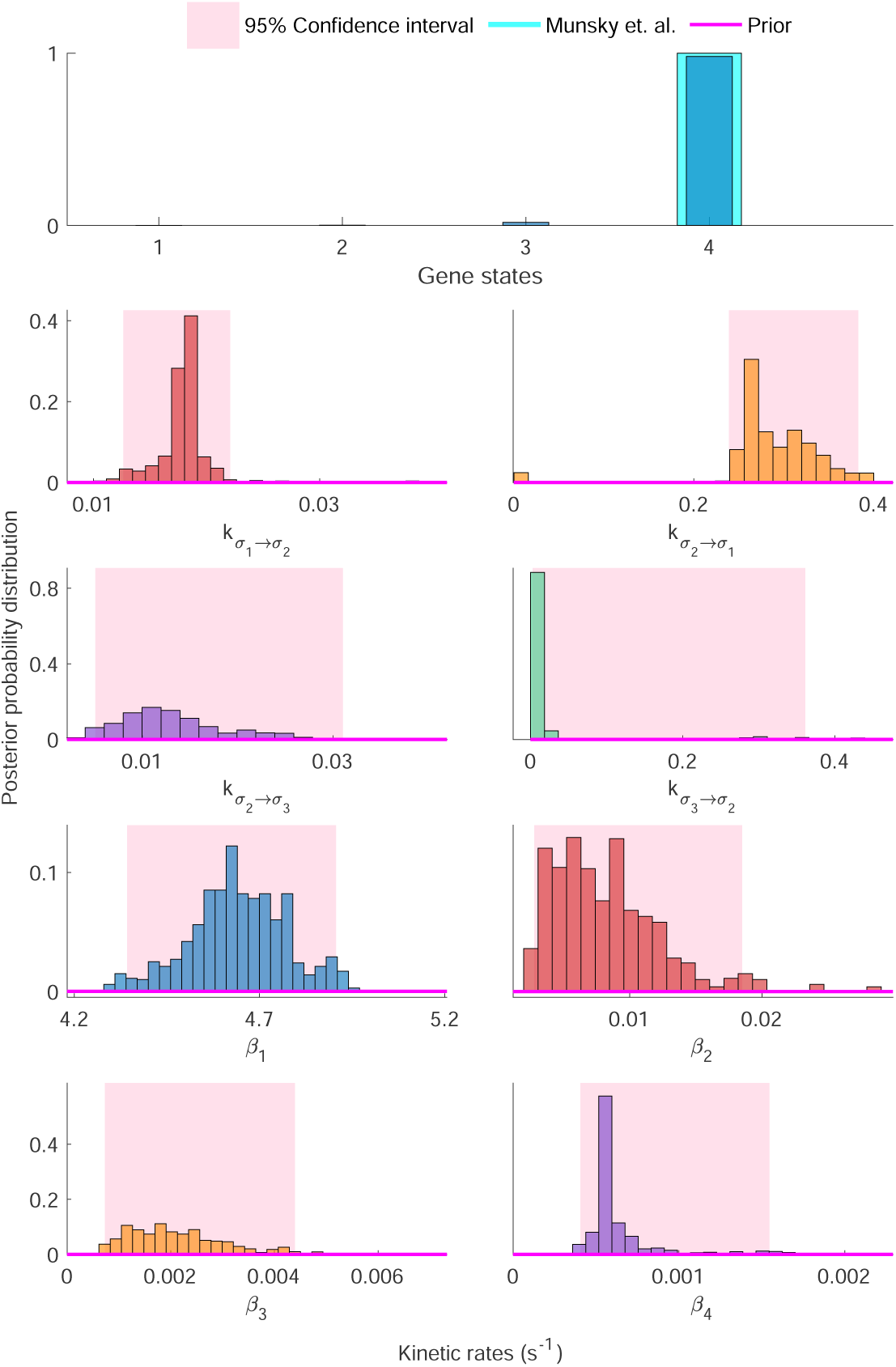
Inference on S. cerevisiae data. Here we show the subset of inferred rates corresponding to linear switching between the four gene states we infer, in agreement with [39] (referenced in the figure as Munsky et. al.). We find that our method infers a transcriptional model with four gene states with high certainty. Additionally, the transition rates correspond closely with so-called ‘linear’ switching (i.e., the gene may leave from each gene state and enter into at most two of the three other gene states). Both these results are in agreement with previously specified models ascribed to chromatin remodeling of the STL1 gene [26]. See Fig. S 11 for joint histograms of rates shown here.

## 4 Methods

### 4.1 Model Formulation

In each gene state, indexed *σ*_*l*_, the gene transcribes RNA copies at rate *β*_*l*_. All RNA degrade stochastically according to overall rate *γ*. A gene can transition stochastically, say from state *σ*_*l*_ to *σ*_*l*_^*′*^, at rate 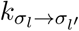. For convenience, we recruit all parameters collectively under the symbol

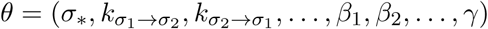

where *σ*_*l*_ = *σ*\* denotes the initial gene state.

In order to infer *θ* within the Bayesian paradigm, we must first specify a likelihood,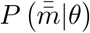. Given measurements 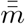, the likelihood is given by

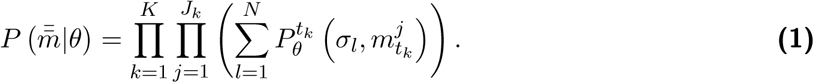

with 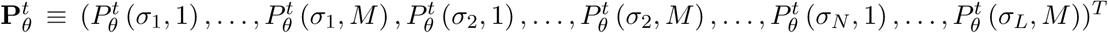 satisfying the CME, 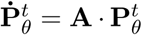, where **A** is a generator matrix, whose dependency on *θ* is outlined in more detail in Section S 1.2.

### 4.2 Model Inference

In order to construct a posterior using our likelihood, we require priors over all model parameters. The priors over quantities in *θ* are chosen for computational convenience alone and are detailed in Section S 1.4.

Here we only expand upon the nonparametric prior used on gene states.

Within a nonparametric formulation, we must theoretically consider an infinite number of gene states and allow the data to winnow down these infinite possibilities to those warranted by the data. This is similar to regular (parametric) Bayesian methods which typically assume broad priors over parameters and eventually allow the data, incorporated through the likelihood, to sharpen parameter estimates (i.e., sharpen the posterior).

As a matter of computational convenience alone, we use the Beta-Bernoulli process [58, 59] as a formal prior on the existence or non-existence of these states.

Put simply, we introduce an infinite number of intermediate binary (Bernoulli) indicator variables, *b*_*l*_, termed loads, which equal 1 when the gene state *σ*_*l*_ is deemed necessary by the data, or 0 otherwise. To make computation feasible, we introduce a so-called weak limit *L*, setting an upper bound on the number of possible gene states [58, 59]. We refer to all loads {*b*_*l*_}_*l*=1:*L*_ collectively as 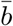.

The Beta-Bernoulli process prior [58, 59] on the loads reads:

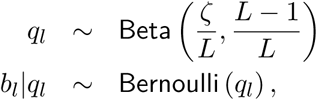

where *q*_*l*_ are hyperparameters describing the success probability of load *b*_*l*_ being “active” or equal to 1, and *ζ* is a hyperhyperparameter. Given this prior, we can learn from the data which gene states are warranted.

Given the likelihood and all priors, we can now construct an explicit form for our posterior probability distribution, 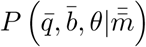. As our likelihood does not assume an analytic form, we generate pseudorandom numbers from 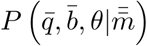 using a custom Markov Chain Monte Carlo (MCMC) [60–62] sampling scheme.

Importantly, it is the ability to efficiently explore this posterior, especially given the added difficulty of inferring states, that will allow us to escape the traps (local maxima) that have impacted the assessment of parameters of other methods Section 3.2.2.

With this fact in mind, we use an overall Gibbs sampling scheme to construct our Markov Chain. Within this Gibbs sampling scheme, we can sample the initial condition, *σ*_*\**_, and loads, 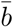, directly from their joint marginal posterior distribution. By contrast, success probabilities 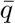 are sampled using a Metropolis-Hastings sampling scheme. Owing to the fact that: 1) we are simultaneously learning discrete (number of gene states, initial condition), and continuous (kinetic rates) parameters; and 2) there is a significant scale separation between various individual continuous parameters, we may encounter featureless posterior distributions over large portions of the possible model space.

To address problem 1), we sample all parameters with Parallel Tempering (PT) in order to better explore the discrete parameters. Within our PT scheme, we propose continuous parameters using Hamiltonian Monte Carlo (HMC) sampling, solving problem 2). Used in conjunction for the first time, these sampling schemes permit the inference of gene states and their associated parameters on reasonable time scales, avoiding local maxima mentioned earlier in Section 3.2.1.

## 5 Discussion

Inferring the most probable regulatory network structure for a given set of observed snapshot RNA expression data presents unique challenges that have stood in the way of accurately identifying the number and connectivity of biophysical reactions and constituent parameters, whether separately or simultaneously. Our approach achieves model and parameter inference in a self-consistent and simultaneous fashion and improves upon limitations of other approaches, including 1) the assumption of steady-state dynamics [63] and 2) separation of model selection of gene state numbers and parameter inference [22, 23, 34, 35].

We evaluated our method’s effectiveness using both experimental and simulated snapshot RNA expression data. For *E. Coli* in fast-growth media, our approach determined (Fig. 5) that a three-state model was more probable compared to the previously utilized two-state model. The additional state is an intermediate production state of the *lacZ* gene, with an intermediate rate of production lying between ‘on’ and ‘off’ rates assumed in a previous analysis [49]. For the *STL1* pathway in *S. cerevisiae*, our approach confirmed that the previously utilized four-state network, with multiple production states, is the most likely model [26, 39]. Critically, the approach detailed here does not *a priori* assume the number or connectivity of gene states. Finally, we demonstrated the robustness of our approach using synthetic snapshot RNA expression data created by simulated regulatory networks designed to challenge any computational inference approach. These results demonstrate a general, simultaneous, self-consistent method to infer gene regulatory models and associated rates from snapshot RNA expression data obtained by smFISH.

We can make several additional extensions to our framework. First, with minimal modification, our approach may utilize spatial information contained in snapshot RNA expression data quantified by smFISH to determine, say, transport rates of RNA from nucleus to cytoplasm. Indeed, the additional constraint of RNA transport from the nucleus to cytoplasm has previously improved parameter identification [39, 64, 65]. Second, modifications of the measurement model within our framework may allow for time-varying rates of transcription, gene state transitions, and RNA degradation [66]. Finally, as the density of gene species increases using highly multiplexed smFISH methods, the flexible network connectivity of our approach may allow regulatory models to explore the most-likely regulatory networks for co-varying gene expression [67–69].

The above generalizations will introduce additional complexity to our likelihood’s computation, already the most costly inference step. The additional complexity is directly due to the increase in the state number and complexity of the connectivity map. Both alter the generator matrix **A** (see Section S 1.2), making it either larger for number of states or denser for connectivity. In the case where the generator matrix remains sparse, the CME solution’s time cost scales roughly linearly with **A**’s size, and FSP based Krylov subspace methods [70, 71] may be more optimal than the CME solution method used here. How the computational cost of **A**’s CME scales with density is more complex than the number of states. Above a certain density, the recently proposed Quantized Tensor Train method [72] may be more efficient, as the FSP based Krylov subspace approach uses incremental time stepping rather than jumping immediately to the times desired for analysis. Alternatively, there have been promising attempts to solve ODEs using neural networks [73]. In addition to facilitating the difficulties arising due to dense CME generator matrices, neural network approaches may further enable parameter inference for non-Markovian models of gene transcription [74].

It may also be possible to deduce gene networks from direct image gene expression dynamics in real-time within living cells [75–79]. However, such approaches obtain real-time kinetics of a limited number of molecules at the expense of higher data density. What is more, genetic manipulation limits the accessible insight to local molecular and biophysical interactions.

The desire to understand downstream consequences of gene expression, for example, through predictive modeling, directly motivates the use of snapshot RNA expression data especially for higher data density. While removing temporal correlations in snapshot RNA expression data immediately hinders the ability to obtain direct insight into gene regulatory dynamics, we may fill the knowledge gaps by increasing the density of time points, RNA species, cells, and stimuli conditions in snapshot RNA expression data. Indeed, here we show how to maximize the information deduced from snapshot RNA expression data and obtain probabilities over gene regulatory network structures and constituent rates. We achieve this by reframing the gene regulatory network identification problem within the Bayesian nonparametric paradigm and developing the requisite tools for inference over gene states, connectivity, and parameters. The probabilistic output of the approach introduced may now allow us to learn networks reflecting the confidence that any given snapshot RNA expression data set supports.

## 6 Acknowledgments

We thank Prof. Ido Golding for providing the experimental data analyzed herein. We thank Prof. Ioannis Sgouralis and Dr. Zachary Fox for interesting discussions and insights. D.P.S. acknowledges support from the NIH (R01HL068702), and S.P. acknowledges support from NIH NIGMS (R01GM130745) and NIH NIGMS (R01GM134426).

