## supplementary information and figures for "Inferring gene expression models from snapshot RNA data"

##### S 1.1 Model overview

Here we outline and describe our computational structure including our measurement model's formulation and the inference strategy's construction. Our method operates on single cell RNA count data, with $m^j_{t_k}$ representing a single cell RNA count for cell $j$ taken at time point $t_k$ for $k = 1, 2, \ldots, K$ and $j = 1, \ldots, J_k$, collected as $\bar{\bar{m}} = \{\bar{m}_{t_k}\}_{k=1}^K$, where $\bar{m}_k = \{m^j_{t_k}\}_{j=1}^J$. Given these single cell RNA counts, we extract information about the underlying gene state model driving the observed dynamics. The gene state model consists of kinetic parameters including transition rates between gene states (and thus gene state connectivity), production rates, and degradation rates.

To infer these kinetic parameters, we must construct a gene state model. The defining characteristic of the model, the number of gene states, ultimately dictates the number of kinetic parameters involved in the underlying dynamics.

Below, we first describe a simpler analog of our method for deriving kinetic parameter approximations when the number of gene states is held at a fixed quantity, i.e., a parametric framework. We then expand our inference framework to the nonparametric paradigm to infer both the number of gene states and kinetic parameters.

##### S 1.2 Model description

Working within a Bayesian paradigm, we must first connect the given single cell RNA counts to the model parameters. For ease of explanation, we start by describing a parametric formulation, for which we will provisionally fix the number of gene states to two. For the two state model, we label each state using $\sigma_l$ for $l = 1, 2$. We can now describe all associated kinetic parameters which include the initial gene state $\sigma_*$, either $\sigma_1$ or $\sigma_2$; the transition rates, $k_{\sigma_1 \to \sigma_2}$ and $k_{\sigma_2 \to \sigma_1}$; the production rates, $\beta_1$ and $\beta_2$; and the degradation rate, $\gamma$, all grouped into $\theta = (\sigma_*, k_{\sigma_1 \to \sigma_2}, k_{\sigma_2 \to \sigma_1}, \beta_1, \beta_2, \gamma)$.

Armed with $\theta$, we relate the kinetic parameters to the single cell RNA counts $\bar{\bar{m}}$ with the measurement model. This measurement model employs the chemical master equation (CME). More specifically, we use the solution to the CME, $P^t_\theta(\sigma_l, m)$, which gives the probability of encountering $m$ RNA in a cell residing in gene state $\sigma_l$ at time $t$ given model parameters $\theta$. Dynamics of these probabilities are governed by a generator matrix, denoted by $\mathbf{A}$, whose structure is dictated by the underlying gene state model.

###### S 1.2.1 Generator Matrix

Here $\mathbf{A}$ is a square block matrix which depends on $\theta$ and whose size reflects the number of gene states.

To describe the structure of $\mathbf{A}$, we start with the nonzero elements associated with the $n$th row, arbitrarily chosen, for which $n < M$ and is therefore associated to the derivative $dP^t_\theta(\sigma_1, n)/dt$. The nonzero elements include

$$\mathbf{A}\left(n,1:2*(M+1)\right) = \begin{Bmatrix} A(n,n-1) \\ A(n,n) \\ A(n,n+1) \\ A(M+1+n,n-1) \end{Bmatrix} = \begin{Bmatrix} \beta_1 \\ -\left(k_{\sigma_1\to\sigma_2}+\beta_1+n\gamma\right) \\ (n+1)\,\gamma \\ k_{\sigma_2\to\sigma_1} \end{Bmatrix}.$$

We note that, as the number of RNA is unbounded, we follow current literature [8] in truncating the infinite set of CME ordinary differential equations (ODEs) by a preset maximum $(M)$ RNA count within a given dataset. Further details on the composition of $\mathbf{A}$ can be found in [42].

Given the the generator matrix's form, $\mathbf{A}$, we define the CME's structure for which the solution, $P^t_\theta(\sigma_l, m)$, satisfies

$$\frac{d}{dt}\mathbf{P}\left(t|\theta\right) = \mathbf{A}\cdot\mathbf{P}\left(t|\theta\right)$$

where

$$\mathbf{P}\left(t|\theta\right) = \Big(P^t_\theta\left(\sigma_1,0\right), P^t_\theta\left(\sigma_1,1\right), \ldots, P^t_\theta\left(\sigma_1,M\right), P^t_\theta\left(\sigma_2,0\right), P^t_\theta\left(\sigma_2,2\right), \ldots, P^t_\theta\left(\sigma_2,M\right)\Big)^T.$$

Provided the above, we now construct our likelihood, $P\left(\bar{\bar{m}}|\theta\right)$, which is a measure of how probable the data, $\bar{\bar{m}}$ is given a set of parameters $\theta$. Sine the single cell RNA counts are independent and identically distributed we build a likelihood of the form

$$P\left(\bar{\bar{m}}|\theta\right) = \prod_{k=1}^{K}\prod_{j=1}^{J_k}\left(\sum_{l=1}^{N} P^{t_k}_\theta\left(\sigma_l, m^j_{t_k}\right)\right). \quad \textbf{(2)}$$

Likelihoods for the one state and three state models follow a similar structure and are described in detail below.

Moving into the nonparametric regime, we must adjust our likelihood dynamically based upon the number of gene states introduced into the gene state model during model inference. Therefore we introduce intermediate binary indicator variables, $b_l$, termed as loads, which correspond to each possible gene state and can be either 1 or 0. The loads affect the configuration of $\mathbf{A}$, and thus the CME's solution, by multiplying all rates associated to their respective gene state. Specifically, the production rate for gene state $n$, $\beta_l$, is multiplied by load $b_l$. Similarly the transition rate from gene state $l$ to gene state $l'$, $k_{\sigma_l\to\sigma'_l}$, is multiplied by loads $b_l$ and $b'_l$ as both gene states must exist for a transition between them to be relevant.

Finally, the degradation rate is multiplied by each respective load, since if a gene state is inactive, the gene will never occupy that state, and therefore degradation will not occur in that state. We demonstrate this change using the same nonzero elements of $\mathbf{A}$ as were outlined for the parametric two state model,

$$\mathbf{A}\left(n, 1: L*(M+1)\right) = \left\{\begin{matrix} A(n,n-1) \\ A(n,n) \\ A(n,n+1) \\ A(M+1+n,n-1) \end{matrix}\right\} = \left\{\begin{matrix} b_1\beta_1 \\ -\left(b_1\sum_{l=2}^{L} b_l k_{\sigma_1\to\sigma_l} + b_1\beta_1 + nb_1\gamma\right) \\ (n+1)\, b_1\gamma \\ b_1\sum_{l=2}^{L} k_{\sigma_l\to\sigma_1} \end{matrix}\right\},$$

where $L$ is chosen as a weak limit imposed on the maximum possible number of gene states for computational feasibility. In this case, this row corresponds to the derivative $dP^t_{\bar{b},\theta}\left(\sigma_1, n\right)/dt$ which is as described for the two state model but now incorporates the new dependencies on the load variables, grouped as $\bar{b} = (b_1, b_2, \ldots, b_L)$, as shown for $\mathbf{A}\left(n, 1: L*(M+1)\right)$.

In this way, each load determines whether or not its corresponding gene state, and thus the associated rates, is warranted by the data. The model parameters encompassed in $\theta$ grow to include all possible transition rates and production rates.

#### S 1.3 Model inference

In the Bayesian paradigm, once we have defined our likelihood, we must place priors over all parameters we wish to infer. Choices for priors over the kinetic parameters as well as the initial condition gene state are relatively straightforward and are described in Section S 1.4. To place priors over the loads, $b_n$, we use Beta-Bernoulli process priors

$$\begin{aligned} q_l &\sim \text{Beta}\left(\frac{\zeta}{L}, \frac{L-1}{L}\right) \\ b_l|q_l &\sim \text{Bernoulli}\left(q_l\right), \end{aligned}$$

where $q_n$ re hyperparameters which describe the success probability of load $b_n$ being "active" or equal to 1, and $\zeta$ is a hyperhyperparameter. Given this setup, we can determine which gene states are deemed necessary by the data.

Given the likelihood and all priors, we can now define our fully joint posterior probability distribution for both parametric, $P\left(\theta|\bar{\bar{m}}\right)$ and nonparametric, $P\left(\bar{q}, \bar{b}, \theta|\bar{\bar{m}}\right)$, frameworks. As there is no possible fully conjugate prior (over all variables) associated to our likelihood, which does not have a closed form solution due to the CME, our posterior probability also does not attain a closed form. Therefore, we must develop a Markov Chain Monte Carlo (MCMC) scheme [60–62] which will allow us to generate pseudo-random samples from our posteriors.

The main bottleneck in our model is the high computational expense associated with numerically approximating CME solutions. In particular, as the size of the CME is given by the structure of the generator matrix, $\mathbf{A}$, the computational cost grows for data sets associated with high RNA counts (highly expressed genes) and gene state models incorporating higher numbers of gene states.

##### S 1.4 Summary of Equations

We define the loads, $\bar{b} = (b_1, b_2, \ldots, b_L)$, transition rates, $\bar{k} = \left(k_{\sigma_l \to \sigma_{l'}}\right)$ for $l = 1, \ldots, L$, $l' = 1, \ldots, L$ and $l \neq l'$, and production rates $\bar{\beta} = (\beta_1, \beta_2, \ldots, \beta_L)$. The success probability vector, $\bar{q} = (q_1, q_2, \ldots, q_L)$ and the initial condition gene state $\sigma_*$. We define, $\tilde{\beta}_l = \log_{10}(\beta_l)$ and keep a similar convention for all other rates. Finally, we collect the set of parameters in $\tilde{\theta} = \left(\tilde{\bar{k}}, \tilde{\bar{\beta}}, \tilde{\gamma}\right)$. Our full set of equations now follows

$$
\begin{aligned}
q_l &\sim \textbf{Beta}\left(\frac{\zeta}{L}, \frac{L-1}{L}\right), l = 1, \ldots, L \\
\sigma_* | \bar{q} &\sim \textbf{Categorical}\left(\frac{\bar{q}}{\sum_{l=1}^{L} q_l}\right) \\
b_l | q_l, \sigma_* &\sim \textbf{Bernoulli}\left(\delta_{\sigma_* l} + (1 - \delta_{\sigma_* l})\, q_l\right), l = 1, \ldots, L \\
\tilde{k}_{\sigma_l \to \sigma_{l'}} &\sim \textbf{Normal}\left(\phi_{1_{ll'}}, \psi_{1_{ll'}}\right), l = 1, \cdots, L, l' = 1, \cdots, J, l \neq l' \\
\tilde{\beta}_l &\sim \textbf{Normal}\left(\phi_{2_l}, \psi_{2_l}\right), l = 1, \ldots, L, \\
\tilde{\gamma} &\sim \textbf{Normal}\left(\phi_3, \psi_3\right) \\
\bar{\bar{m}} | \sigma_*, \bar{b}, \theta &\sim \prod_{k=1}^{K} \prod_{j=1}^{J_k} \left(\sum_{i=1}^{L} P^{t, \sigma_*, \bar{b}, \theta}\left(\sigma_i, m_k^j\right)\right).
\end{aligned}
$$

##### S 1.5 Nonparametric Chemical Master Equation

Let $P^{t,\sigma_*,\bar{b},\theta}(\sigma_l, m)$ be the probability of a cell being in gene state $\sigma_l$ with $m$ RNA count, $m$, at time $t$ given an initial gene state of $\sigma_*$ and parameters $\theta$, such that

$$
\bar{\mathbf{P}}^{t,\sigma_*,\bar{b},\theta}(\sigma_l, m) = \left(P^{t,\sigma_*,\bar{b},\theta}(\sigma_l, 0), P^{t,\sigma_*,\bar{b},\theta}(\sigma_l, 1), \ldots, P^{t,\sigma_*,\bar{b},\theta}(\sigma_l . M)\right),
$$

where $M$ is the maximum number of RNA per cell considered (and fixed at some high albeit reasonable number for computational efficiency). Then the master equation for the BNP model, in matrix form, reads

$$
\frac{d}{dt}\left(\begin{array}{c}\bar{\mathbf{P}}^{t,\sigma_*,\bar{b},\theta}(\sigma_1,m)\\ \hline \bar{\mathbf{P}}^{t,\sigma_*,\bar{b},\theta}(\sigma_2,m)\\ \hline \vdots \\ \hline \bar{\mathbf{P}}^{t,\sigma_*,\bar{b},\theta}(\sigma_L,m)\end{array}\right) = \left(\begin{array}{c|c|c|c}\mathbf{T}_1 & \mathbf{K}_{\sigma_2\to\sigma_1} & \cdots & \mathbf{K}_{\sigma_L\to\sigma_1}\\ \hline \mathbf{K}_{\sigma_1\to\sigma_2} & \mathbf{T}_2 & \ddots & \mathbf{K}_{\sigma_L\to\sigma_2}\\ \hline \vdots & \ddots & \ddots & \vdots \\ \hline \mathbf{K}_{\sigma_1\to\sigma_L} & \mathbf{K}_{\sigma_2\to\sigma_L} & \cdots & \mathbf{T}_L\end{array}\right)\left(\begin{array}{c}\bar{\mathbf{P}}^{t,\sigma_*,\bar{b},\theta}(\sigma_1,m)\\ \hline \bar{\mathbf{P}}^{t,\sigma_*,\bar{b},\theta}(\sigma_2,m)\\ \hline \vdots \\ \hline \bar{\mathbf{P}}^{t,\sigma_*,\bar{b},\theta}(\sigma_L,m)\end{array}\right)
$$

with the matrix,

$$
\mathbf{T}_l = \begin{pmatrix}
-\left(b_l\beta_l + \sum_{j=1}^{L} b_l b_{l'} k_{\sigma_l\to\sigma_{l'}}\right) & b_l\gamma & 0 & 0 & \cdots \\
b_l\beta_l & -\left(b_l\beta_l + \sum_{l'=1}^{L} b_l b_{l'} k_{\sigma_l\to\sigma_{l'}} + b_l\gamma\right) & 2b_l\gamma & 0 & \ddots \\
0 & b_l\beta_l & -\left(b_l\beta_l + \sum_{j=1}^{L} b_l b_{l'} k_{\sigma_l\to\sigma_{l'}} + 2b_l\gamma\right) & 3b_l\gamma & \ddots \\
\vdots & \ddots & \ddots & \ddots & \ddots \\
0 & 0 & \cdots & b_l\beta_l & -\left(\sum_{l'=1}^{L} b_l b_{l'} k_{\sigma_l\to\sigma_l'} + Mb_l\gamma\right)
\end{pmatrix},
$$

provided that

$k_{\sigma_l\to\sigma_{l'}} = 0$ for $l = 1,\ldots,L$, and $\mathbf{K}_{\sigma_1\to\sigma_{1'}}$ are diagonal matrices with $k_{\sigma_l\to\sigma_{l'}}$ along the diagonals for $l = 1,\ldots,L$, and $l' = 1,\ldots,L$.

We use initial condition

$$
\begin{aligned}
P^{0,\sigma_*,\bar{b},\theta}(\sigma_l,m) &= 1, \quad \text{for } l = \sigma_* \text{ and } m = 0 \\
P^{0,\sigma_*,\bar{b},\theta}(\sigma_l,m) &= 0, \quad \text{otherwise.}
\end{aligned}
$$

##### S 1.6 Description of the Computational Scheme

We use an overall Gibbs sampling scheme to construct our Markov Chain. Clearly, the Gibbs scheme is a method for updating a the full set of parameters which completely parameterize a model. Let's call this complete set of parameters $\Lambda$. In each step of the Gibbs scheme, we obtain a new set of parameters, $\Lambda$, from a set of old parameters $\Lambda^{old}$ by proposing multiple disjoint subsets $\theta_i^{prop}$ of $\Lambda$ for $i = 1 : I$, which together, would completely parameterize the model. For each $i$ in sequence, we provisionally update the full set of parameters to $\Lambda^{prop}$, using the values which make up $\theta_i^{prop}$ and compare the posterior probability of $\Lambda^{prop}$ (which now contains the parameters from $\theta_i^{prop}$ in addition to the most recent remaining parameters in $\Lambda$), to that of $\Lambda$. If the comparison favors $\Lambda^{prop}$, we make the update

$\Lambda = \Lambda^{prop}$, otherwise $\Lambda$ remains unchanged. In this way, once all $I$ subsets have been assessed and either accepted or rejected, we obtain a new estimates for all parameters in $\Lambda$.

Within our Gibbs sampling scheme, we sample the initial condition, $\sigma_*$ and $\bar{b}$ directly from their joint marginal posterior distribution (Section S 1.6.1). By contrast, success probabilities $\bar{q}$ are sampled using a Metropolis-Hastings sampling scheme (Section S 1.6.2). Owing to the fact that: 1) we are simultaneously learning discrete (number of gene states, initial condition), and continuous (kinetic rates) parameters; and 2) there is a significant scale separation between various individual continuous parameters, we may encounter featureless posterior distributions over large portions of the possible model space. To address problem 1), we sample all parameters with Parallel Tempering (PT) in order to better explore the discrete parameters. Within our PT scheme, we propose all continuous parameters (kinetic rates) using Hamiltonian Monte Carlo (HMC) (Section S 1.6.6) sampling, solving problem 2).

##### S 1.6.1 Joint sampling of the loads and initial condition gene state

We use direct sampling within a Gibbs sampling scheme to sample the initial condition, $\sigma_*$ and $b_l$ for $l = 1, \dots, L$, jointly from the marginal posterior distribution

$$
\begin{aligned}
\mathbb{P}\left(\sigma_*, \bar{b} | \bar{\bar{m}}, \bar{q}, \theta\right) &\propto \mathbb{P}\left(\bar{\bar{m}} | \sigma_*, \bar{b}, \theta\right) \mathbb{P}\left(\bar{b} | \bar{q}, \sigma_*\right) \mathbb{P}\left(\sigma_* | \bar{q}\right) \\
&= \prod_{k=1}^{K} \prod_{j=1}^{J_k} \left( \sum_{l=1}^{L} P^{t,\sigma_*,\bar{b},\theta}\left(\sigma_l, m_k^j\right)\right) \prod_{l}^{L} \textbf{Bernoulli}\left(b_l; \delta_{\sigma_* l} + \left(1 - \delta_{\sigma_* l}\right) q_l\right) \\
&\times \textbf{Categorical}\left(\sigma_*; \frac{\bar{q}}{\sum\limits_{l=1}^{L} q_l}\right) \\
&= \prod_{k=1}^{K} \prod_{j=1}^{J_k} \left( \sum_{l=1}^{L} P^{t,\sigma_*,\bar{b},\theta}\left(\sigma_l, m_k^j\right)\right) \\
&\times \prod_{l}^{L} \left(\delta_{\sigma_* l} + \left(1 - \delta_{\sigma_* l}\right) q_l\right)^{b_l} \left(1 - \left(\delta_{\sigma_* l} + \left(1 - \delta_{\sigma_* l}\right) q_l\right)\right)^{1 - b_l} \left(\frac{q_{\sigma_*}}{\sum\limits_{l=1}^{L} q_l}\right).
\end{aligned}
$$

We will sample the initial condition gene state and the loads through posteriors of all possible initial condition gene states with all possible load combinations

$$
\begin{aligned}
P_1 &= \mathbb{P}\left(C_1 | \bar{\bar{m}}, \bar{\phi}_*, \theta\right), & C_1 &= \left(\sigma_* = 1, \bar{b} = [1, 0, \dots, 0]\right) \\
P_2 &= \mathbb{P}\left(C_2 | \bar{\bar{m}}, \bar{\phi}_*, \theta\right), & C_2 &= \left(\sigma_* = 1, \bar{b} = [1, 1, \dots, 0]\right) \\
&\vdots \\
P_{2^{L-1}} &= \mathbb{P}\left(C_{2^{L-1}} | \bar{\bar{m}}, \bar{\phi}_*, \theta\right), & C_{2^{L-1}} &= \left(\sigma_* = 1, \bar{b} = [1, 1, \dots, 1]\right)
\end{aligned}
$$

$$P_{2^{L-1}+1} = \mathbb{P}\left(C_{2^{L-1}+1}|\bar{\bar{m}}, \bar{\phi}_*, \theta\right), \quad C_{2^{L-1}+1} = \left(\sigma_* = 2, \bar{b} = [0, 1, \ldots, 0]\right)$$
$$\vdots$$
$$P_{2*2^{L-1}} = \mathbb{P}\left(C_{2*2^{L-1}}|\bar{\bar{m}}, \bar{\phi}_*, \theta\right), \quad C_{2*2^{L-1}} = \left(\sigma_* = 2, \bar{b} = [1, 1, \ldots, 1]\right)$$
$$\vdots$$
$$P_{L*2^{L-1}} = \mathbb{P}\left(C_{L*2^{L-1}}|\bar{\bar{m}}, \bar{\phi}_*, \theta\right), \quad C_{L*2^{L-1}} = \left(\sigma_* = L, \bar{b} = [1, 1, \ldots, 1]\right).$$

and construct the Categorical distribution and sample the initial condition gene state

$$\sigma_*, \bar{b}|\bar{\bar{m}}, \bar{q}, \theta \quad \sim \quad \textbf{Categorical}_{[C_1, C_2, \ldots, C_{L*2^{L-1}}]}\left(P_1, P_2, \ldots, P_{L*2^{L-1}}\right).$$

The conditional probabilities can be written

$$\begin{aligned}
\mathbb{P}\left(\sigma_* = l', \bar{b}|\bar{\bar{m}}, \bar{q}, \theta\right) &\propto \mathbb{P}\left(\bar{\bar{m}}|\sigma_*, \bar{b}, \theta\right)\mathbb{P}\left(\bar{b}|\bar{q}, \sigma_*\right)\mathbb{P}\left(\sigma_*|\bar{q}\right) \\
&= \prod_{k=1}^{K}\prod_{j=1}^{J_k}\left(\sum_{l=1}^{L} P^{t,l',\bar{b},\theta}\left(\sigma_l, m_k^j\right)\right) \\
&\times \prod_{l}^{L}\left(\delta_{l'l} + (1-\delta_{l'l})\,q_l\right)^{b_l}\left(1 - \left(\delta_{l'l} + (1-\delta_{l'l})\,q_l\right)\right)^{1-b_l}\left(\frac{q_{l'}}{\sum\limits_{l=1}^{L} q_l}\right)
\end{aligned}$$

The logarithm of the conditional probability

$$\begin{aligned}
\log\left(\mathbb{P}\left(\sigma_* = l', \bar{b}|\bar{\bar{m}}, \bar{q}, \theta\right)\right) &= \sum_{k=1}^{K}\sum_{j=1}^{J_k}\log\left(\sum_{l=1}^{L} P^{t,l',\theta}\left(\sigma_l, m_k^j\right)\right) \\
&+ \sum_{l=1}^{L}\left(b_l\left(\log\left(\delta_{l'l} + (1-\delta_{l'l})\,q_l\right)\right)\right. \\
&+ \left.(1-b_l)\left(\log\left(1 - \left(\delta_{l'l} + (1-\delta_{l'l})\,q_l\right)\right)\right)\right) \\
&+ \log\left(q_{l'}\right) - \log\left(\sum_{l=1}^{L} q_l\right).
\end{aligned}$$

We use a mixture of MH and Hamiltonian Monte Carlo (HMC) sampling schemes to sample the reaction rates. In both cases we sample from the marginal posterior distribution

$$\mathbb{P}\left(\tilde{\theta}|\bar{\bar{m}}, \bar{q}, \sigma_*, \bar{b}\right) \quad \propto \quad \mathbb{P}\left(\bar{\bar{m}}|\sigma_*, \bar{b}, \tilde{\theta}\right)\mathbb{P}\left(\tilde{\theta}\right).$$

##### S 1.6.2 Sampling Success Probabilities

We use a Metropolis-Hastings (MH) sampling scheme to sample the initial success probability vector, $\bar{q}$ from the marginal posterior distribution

$$\begin{aligned}
\mathbb{P}\left(q_l|\bar{\bar{m}}, \sigma_*, \bar{b}, \theta\right) &\propto \mathbb{P}\left(\bar{b}|\sigma_*, \bar{q}\right)\mathbb{P}\left(\sigma_*|\bar{q}\right)\mathbb{P}\left(\bar{q}\right) \\
&= \prod_{l=1}^{L} \textbf{Bernoulli}\left(b_l; \delta_{\sigma_* l} + (1-\delta_{\sigma_* l})\, q_l\right) \textbf{Categorical}\left(\sigma_*; \frac{\bar{q}}{\sum\limits_{l=1}^{L} q_l}\right) \\
&\times \prod_{l=1}^{L} \textbf{Beta}\left(q_l; \frac{\zeta}{L}, \frac{L-1}{L}\right) \\
&= \prod_{l}^{L} \left(\delta_{\sigma_* l} + (1-\delta_{\sigma_* l})\, q_l\right)^{b_l} \left(1 - \left(\delta_{\sigma_* l} + (1-\delta_{\sigma_* l})\, q_l\right)\right)^{1-b_l} \left(\frac{q_{\sigma_*}}{\sum\limits_{l=1}^{L} q_l}\right) \\
&\times \prod_{l=1}^{L} \frac{1}{B\left(\frac{\zeta}{L}, \frac{L-1}{L}\right)} q_l^{\frac{\zeta}{L}-1} \left(1-q_l\right)^{\frac{L-1}{L}-1}.
\end{aligned}$$

We propose directly from the prior distribution, giving an acceptance ratio of

$$\begin{aligned}
R_{\bar{q}^{old}}\left(\bar{q}^{prop}\right) &= \frac{\mathbb{P}\left(\bar{b}^{old}|\sigma_*^{old}, \bar{q}^{prop}\right)\mathbb{P}\left(\sigma_*^{old}|\bar{q}^{prop}\right)}{\mathbb{P}\left(\bar{b}^{old}|\sigma_*^{old}, \bar{q}^{old}\right)\mathbb{P}\left(\sigma_*^{old}|\bar{q}^{old}\right)} \\
&= \frac{\prod\limits_{l}^{L} \left(\delta_{\sigma_* l} + (1-\delta_{\sigma_* l})\, q_l^{prop}\right)^{b_l^{old}} \left(1 - \left(\delta_{\sigma_* l} + (1-\delta_{\sigma_* l})\, q_l^{prop}\right)\right)^{1-b_l^{old}}}{\prod\limits_{l}^{L} \left(\delta_{\sigma_* l} + (1-\delta_{\sigma_* l})\, q_l^{old}\right)^{b_l^{old}} \left(1 - \left(\delta_{\sigma_* l} + (1-\delta_{\sigma_* l})\, q_l^{old}\right)\right)^{1-b_l^{old}}} \\
&\times \frac{\frac{q_{\sigma_*}^{prop}}{\sum\limits_{l=1}^{L} q_l^{prop}}}{\frac{q_{\sigma_*}^{old}}{\sum\limits_{l=1}^{L} q_l^{old}}}.
\end{aligned}$$

In log form

$$\begin{aligned}
\log\left(R_{\bar{q}^{old}}\left(\bar{q}^{prop}\right)\right) &= \sum_{l}^{L} \left(b_l^{old}\left(\log\left(\delta_{\sigma_* l} + (1-\delta_{\sigma_* l})\, q_l^{prop}\right) - \log\left(\delta_{\sigma_* l} + (1-\delta_{\sigma_* l})\, q_l^{old}\right)\right)\right. \\
&+ \left.\left(1 - b_l^{old}\right)\left(\log\left(1 - \delta_{\sigma_* l} + (1-\delta_{\sigma_* l})\, q_l^{prop}\right) - \log\left(1 - \delta_{\sigma_* l} + (1-\delta_{\sigma_* l})\, q_l^{old}\right)\right)\right) \\
&+ \log\left(q_{\sigma_*}^{prop}\right) - \log\left(\sum_{l=1}^{L} q_l^{prop}\right) - \log\left(q_{\sigma_*}^{old}\right) + \log\left(\sum_{l=1}^{L} q_l^{old}\right).
\end{aligned}$$

##### S 1.6.3 Sampling the initial condition gene state

We use direct sampling within a Gibbs sampling scheme to sample the initial condition, $\sigma_*$ from the marginal posterior distribution

$$\begin{aligned}
\mathbb{P}\left(\sigma_*|\bar{\bar{m}}, \bar{q}, \bar{b}, \theta\right) &\propto \mathbb{P}\left(\bar{\bar{m}}|\sigma_*, \bar{b}, \theta\right) \mathbb{P}\left(\bar{b}|\bar{q}, \sigma_*\right) \mathbb{P}\left(\sigma_*|\bar{q}\right) \\
&= \prod_{k=1}^{K}\prod_{j=1}^{J_k}\left(\sum_{l=1}^{L} P^{t,\sigma_*,\bar{b},\theta}\left(\sigma_l, m_k^j\right)\right)\prod_{l}^{L}\textbf{Bernoulli}\left(b_l; \delta_{\sigma_* l} + \left(1-\delta_{\sigma_* l}\right) q_l\right) \\
&\times \textbf{Categorical}\left(\sigma_*; \frac{\bar{q}}{\sum_{l=1}^{L} q_l}\right) \\
&= \prod_{k=1}^{K}\prod_{j=1}^{J_k}\left(\sum_{l=1}^{L} P^{t,\sigma_*,\bar{b},\theta}\left(\sigma_l, m_k^j\right)\right) \\
&\times \prod_{l}^{L}\left(\delta_{\sigma_* l} + \left(1-\delta_{\sigma_* l}\right) q_l\right)^{b_l}\left(1-\left(\delta_{\sigma_* l} + \left(1-\delta_{\sigma_* l}\right) q_l\right)\right)^{1-b_l}\left(\frac{q_{\sigma_*}}{\sum_{l=1}^{L} q_l}\right).
\end{aligned}$$

We will sample the initial condition gene state through posteriors of all possible gene states

$$\begin{aligned}
\sigma_* = 1, &\quad P_1 = \mathbb{P}\left(\sigma_* = 1|\bar{\bar{m}}, \bar{\phi}_*, \theta\right) \\
\sigma_* = 2, &\quad P_2 = \mathbb{P}\left(\sigma_* = 2|\bar{\bar{m}}, \bar{\phi}_*, \theta\right) \\
&\vdots \\
\sigma_* = L, &\quad P_L = \mathbb{P}\left(\sigma_* = L|\bar{\bar{m}}, \bar{\phi}_*, \theta\right)
\end{aligned}$$

and construct the Categorical distribution and sample the initial condition gene state

$$\sigma_*|\bar{\bar{m}}, \bar{q}, \bar{b}, \theta \quad \sim \quad \textbf{Categorical}_{[\sigma_1,\sigma_2,\ldots,\sigma_L]}\left(P_1, P_2, \ldots, P_L\right).$$

The conditional probabilities can be written

$$\begin{aligned}
\mathbb{P}\left(\sigma_* = l'|\bar{\bar{m}}, \bar{q}, \bar{b}, \theta\right) &\propto \mathbb{P}\left(\bar{\bar{m}}|\sigma_*, \bar{b}, \theta\right) \mathbb{P}\left(\bar{b}|\bar{q}, \sigma_*\right) \mathbb{P}\left(\sigma_*|\bar{q}\right) \\
&= \prod_{k=1}^{K}\prod_{j=1}^{J_k}\left(\sum_{l=1}^{L} P^{t,l',\bar{b},\theta}\left(\sigma_l, m_k^j\right)\right) \\
&\times \prod_{l}^{L}\left(\delta_{l' l} + \left(1-\delta_{l' l}\right) q_l\right)^{b_l}\left(1-\left(\delta_{l' l} + \left(1-\delta_{l' l}\right) q_l\right)\right)^{1-b_l}\left(\frac{q_{l'}}{\sum_{l=1}^{L} q_l}\right).
\end{aligned}$$

The logarithm of the conditional probability

$$\begin{aligned}
\log\left(\mathbb{P}\left(\sigma_* = l'|\bar{\bar{m}}, \bar{q}, \bar{b}, \theta\right)\right) &= \sum_{k=1}^{K}\sum_{j=1}^{J_k}\log\left(\sum_{l=1}^{L} P^{t,l',\theta}\left(\sigma_l, m_k^j\right)\right) \\
&+ \sum_{l=1}^{L}\left(b_l\left(\log\left(\delta_{l'l} + (1-\delta_{l'l})\, q_l\right)\right)\right. \\
&+ \left.(1-b_l)\log\left(1-\delta_{l'l} + (1-(\delta_{l'l})\, q_l)\right)\right) \\
&+ \log\left(q_{l'}\right) - \log\left(\sum_{l=1}^{L} q_l\right).
\end{aligned}$$

##### S 1.6.4 Sampling of the loads

We use direct sampling within a Gibbs sampling scheme to sample the loads, $\bar{b}$ from the marginal posterior distribution

$$\begin{aligned}
\mathbb{P}\left(\bar{b}|\bar{\bar{m}}, \bar{q}, \sigma_*, \bar{k}, \bar{\beta}, \gamma\right) &\propto \mathbb{P}\left(\bar{\bar{m}}|\sigma_*, \bar{b}, \theta\right)\mathbb{P}\left(\bar{b}|\bar{q}, \sigma_*\right) \\
&= \prod_{k=1}^{K}\prod_{j=1}^{J_k}\left(\sum_{l=1}^{L} P^{t,\sigma_*,\bar{b},\theta}\left(\sigma_l, m_k^j\right)\right)\prod_{l}^{L}\textbf{Bernoulli}\left(b_l; \delta_{\sigma_* l} + (1-\delta_{\sigma_* l})\, q_l\right) \\
&= \prod_{k=1}^{K}\prod_{j=1}^{J_k}\left(\sum_{l=1}^{L} P^{t,\sigma_*,\bar{b},\theta}\left(\sigma_l, m_k^j\right)\right) \\
&\times \prod_{l}^{L}\left(\delta_{\sigma_* l} + (1-\delta_{\sigma_* l})\, q_l\right)^{b_l}\left(1-\left(\delta_{\sigma_* l} + (1-\delta_{\sigma_* l})\, q_l\right)\right)^{1-b_l}.
\end{aligned}$$

In order to sample the loads using direct sampling within a Gibbs sampling scheme, we will sample the loads simultaneously through posteriors of all configurations of loads

$$\begin{aligned}
P_1 = \mathbb{P}\left(B_1|\bar{\bar{m}}, \bar{\phi}_*, \sigma_*, \bar{k}, \bar{\beta}, \gamma\right), &\quad B_1 &= [0, 0, \ldots, 0] \\
P_2 = \mathbb{P}\left(B_2|\bar{\bar{m}}, \bar{\phi}_*, \sigma_*, \bar{k}, \bar{\beta}, \gamma\right), &\quad B_2 &= [1, 0, \ldots, 0] \\
&\vdots \\
P_{2^L} = \mathbb{P}\left(B_{2^L}|\bar{\bar{m}}, \bar{\phi}_*, \sigma_*, \bar{k}, \bar{\beta}, \gamma\right), &\quad B_{2^L} &= [1, 1, \ldots, 1]
\end{aligned}$$

and construct the Categorical distribution and sample the configuration of loads

$$\bar{b}|\bar{\bar{m}}, \bar{q}, \sigma_*, \theta \sim \textbf{Categorical}_{\left[B_1, B_2, \ldots, B_{2^L}\right]}\left(P_1, P_2, \ldots, P_{2^L}\right).$$

However, since the Categorical distribution has $2^L$ arguments, we lessen the computational cost by sampling a smaller set of loads. We define a random set of loads $\bar{b}_{l'}$ such that $l' \neq \sigma_*$ for all $l'$ and apply direct sampling to these. The posterior over this smaller set of nodes that we update simultaneously is then $\mathbb{P}\left(\bar{b}_{l'}|\bar{\bar{m}}, \bar{q}, \sigma_*, \bar{b}_{-l'}, \bar{k}, \bar{\beta}, \gamma\right)$ in which $\bar{b}_{-l'}$ is the set $\bar{b}$ excluding the loads contained in $\bar{b}_{l'}$. Thus, the conditional probability can then be written as

$$
\begin{aligned}
\mathbb{P}\left(\bar{b}_{l'}|\bar{\bar{m}}, \bar{q}, \sigma_*, \bar{b}_{-l'}\theta\right) &\propto \mathbb{P}\left(\bar{\bar{m}}|\sigma_*, \bar{b}, \theta\right)\mathbb{P}\left(\bar{b}|\bar{q}, l'\right) \\
&= \prod_{k=1}^{K}\prod_{j=1}^{J_k}\left(\sum_{l=1}^{L} P^{t,\sigma_*,\bar{b},\theta}\left(\sigma_l, m_k^j\right)\right)\prod_{l}^{L}\textbf{Bernoulli}\left(b_l; \delta_{\sigma_* l} + (1-\delta_{\sigma_* l})\, q_l\right) \\
&= \prod_{k=1}^{K}\prod_{j=1}^{J_k}\left(\sum_{l=1}^{L} P^{t,\sigma_*,\bar{b},\theta}\left(\sigma_l, m_k^j\right)\right) \\
&\times \prod_{l}^{L}\left(\delta_{l'l} + (1-\delta_{\sigma_* l})\, q_l\right)^{b_l}\left(1 - \left(\delta_{\sigma_* l} + (1-\delta_{\sigma_* l})\, q_l\right)\right)^{1-b_l}.
\end{aligned}
$$

The logarithm of the conditional probability is

$$
\begin{aligned}
\log\left(\mathbb{P}\left(\bar{b}_{l'}|\bar{\bar{m}}, \bar{q}, \sigma_*, \bar{b}_{-l'}\theta\right)\right) &= \sum_{k=1}^{K}\sum_{j=1}^{J_k}\log\left(\sum_{l=1}^{L} P^{t,\sigma_*,\bar{b}_{l'},\bar{b}_{-l'},\theta}\left(\sigma_l, m_k^j\right)\right) \\
&+ \sum_{l'}\left(b_{l'}\left(\log\left(\delta_{\sigma_* l'} + (1-\delta_{\sigma_* l'})\, q_{l'}\right)\right)\right. \\
&+ \left.(1-b_{l'})\log\left(1 - \left(\delta_{\sigma_* l'} + \left(1 - \left(\delta_{\sigma_* l'}\right) q_{l'}\right)\right)\right)\right).
\end{aligned}
$$

##### S 1.6.5 Metropolis Hastings sampling scheme

We propose new samples using a multivariate normal distribution demonstrated below for the parameters $\tilde{\beta}_l$ and $\tilde{\gamma}$,

$$
\mathbb{Q}_{\tilde{\beta}_l^{old},\tilde{\gamma}^{old}}\left(\tilde{\beta}_l^{prop}, \tilde{\gamma}^{prop}\right) = \textbf{MVNormal}\left(\begin{pmatrix}\tilde{\beta}_l^{prop} \\ \tilde{\gamma}^{prop}\end{pmatrix}; \begin{pmatrix}\tilde{\beta}_l^{old} \\ \tilde{\gamma}^{old}\end{pmatrix}, \mathbf{\Sigma}_{[2_l,3]}\right),
$$

where $\mathbf{\Sigma}_{[2_l,3]}$ is an adapted covariance matrix of size $2 \times 2$ with the corresponding pieces of the larger $N \times N$ covariance matrix $\mathbf{\Sigma}$ for $\beta_l$ and $\gamma$. Continuing with this example, the acceptance ratio would then be

$$
\begin{aligned}
R_{\tilde{\beta}_l^{old},\tilde{\gamma}^{old}}\left(\tilde{\beta}_l^{prop}, \tilde{\gamma}^{prop}\right) &= \frac{\mathbb{P}\left(\bar{\bar{m}}|l_*^{old}, \bar{b}^{old}, \tilde{\theta}^{prop}\right)\mathbb{P}\left(\tilde{\theta}^{prop}\right)}{\mathbb{P}\left(\bar{\bar{m}}|\sigma_*^{old}, \bar{b}^{old}, \tilde{\theta}^{old}\right)\mathbb{P}\left(\tilde{\theta}^{old}\right)} \\
&= \frac{\prod_{k=1}^{K}\prod_{j=1}^{J_k}\left(\sum_{i=1}^{L} P^{t,\sigma_*^{old},\bar{b}^{old},10^{\tilde{\theta}^{prop}}}\left(\sigma_i, m_k^j\right)\right)}{\prod_{k=1}^{K}\prod_{j=1}^{J_k}\left(\sum_{i=1}^{L} P^{t,\sigma_*^{old},\bar{b}^{old},10^{\tilde{\theta}^{old}}}\left(\sigma_i, m_k^j\right)\right)} \\
&\times \frac{\textbf{Normal}\left(\tilde{\beta}_l^{prop}; \phi_{2_l}, \psi_{2_l}\right)\textbf{Normal}\left(\tilde{\gamma}^{prop}; \phi_3, \psi_3\right)}{\textbf{Normal}\left(\tilde{\beta}_l^{old}; \phi_{2_l}, \psi_{2_l}\right)\textbf{Normal}\left(\tilde{\gamma}^{old}; \phi_3, \psi_3\right)}
\end{aligned}
$$

$$
\begin{aligned}
\times & \quad \frac{\mathbb{Q}_{\tilde{\beta}_l^{prop},\tilde{\gamma}^{prop}}\left(\tilde{\beta}_l^{old},\tilde{\gamma}^{old}\right)}{\mathbb{Q}_{\tilde{\beta}_l^{old},\tilde{\gamma}^{old}}\left(\tilde{\beta}_l^{prop},\tilde{\gamma}^{prop}\right)} \\
= & \quad \frac{\prod_{k=1}^{K}\prod_{j=1}^{J_k}\left(\sum_{i=1}^{L} P^{t,\sigma_*^{old},\bar{b}^{old},10^{\tilde{\theta}^{prop}}}\left(\sigma_i, m_k^j\right)\right)}{\prod_{k=1}^{K}\prod_{j=1}^{J_k}\left(\sum_{i=1}^{L} P^{t,\sigma_*^{old},\bar{b}^{old},10^{\tilde{\theta}^{old}}}\left(\sigma_i, m_k^j\right)\right)} \\
\times & \quad \frac{e^{-\frac{1}{2}\left(\frac{\tilde{\beta}_l^{prop}-\phi_{2_l}}{\psi_{2_l}}\right)^2} e^{-\frac{1}{2}\left(\frac{\tilde{\gamma}^{prop}-\phi_3}{\psi_3}\right)^2}}{e^{-\frac{1}{2}\left(\frac{\tilde{\beta}_l^{old}-\phi_{2_l}}{\psi_{2_l}}\right)^2} e^{-\frac{1}{2}\left(\frac{\tilde{\gamma}^{old}-\phi_3}{\psi_3}\right)^2}} \\
\times & \quad \frac{e^{-\frac{1}{2}\left(\begin{bmatrix}\tilde{\beta}_l^{old}\\ \tilde{\gamma}^{old}\end{bmatrix}-\begin{bmatrix}\tilde{\beta}_l^{prop}\\ \tilde{\gamma}^{prop}\end{bmatrix}\right)^T \mathbf{\Sigma}_{[2_l,3]}^{-1}\left(\begin{bmatrix}\tilde{\beta}_l^{old}\\ \tilde{\gamma}^{old}\end{bmatrix}-\begin{bmatrix}\tilde{\beta}_l^{prop}\\ \tilde{\gamma}^{prop}\end{bmatrix}\right)}}{e^{-\frac{1}{2}\left(\begin{bmatrix}\tilde{\beta}_l^{prop}\\ \tilde{\gamma}^{prop}\end{bmatrix}-\begin{bmatrix}\tilde{\beta}_l^{old}\\ \tilde{\gamma}^{old}\end{bmatrix}\right)^T \mathbf{\Sigma}_{[2_l,3]}^{-1}\left(\begin{bmatrix}\tilde{\beta}_l^{old}\\ \tilde{\gamma}^{old}\end{bmatrix}-\begin{bmatrix}\tilde{\beta}_l^{prop}\\ \tilde{\gamma}^{prop}\end{bmatrix}\right)}}.
\end{aligned}
$$

In log form this becomes

$$
\begin{aligned}
\log\left(R_{\tilde{\beta}_l^{old},\tilde{\gamma}^{old}}\left(\tilde{\beta}_l^{prop},\tilde{\gamma}^{prop}\right)\right) = & \quad \sum_{k=1}^{K}\sum_{j=1}^{J_k}\log\left(\sum_{i=1}^{L} P^{t,\sigma_*^{old},\bar{b}^{old},10^{\tilde{\theta}^{prop}}}\left(\sigma_i, m_k^j\right)\right) \\
- & \quad \sum_{k=1}^{K}\sum_{j=1}^{J_k}\log\left(\sum_{i=1}^{L} P^{t,\sigma_*^{old},\bar{b}^{old},10^{\tilde{\theta}^{old}}}\left(\sigma_i, m_k^j\right)\right) \\
- & \quad \frac{1}{2}\left(\frac{\tilde{\beta}_l^{prop}-\phi_{2_l}}{\psi_{2_l}}\right)^2 - \frac{1}{2}\left(\frac{\tilde{\gamma}^{prop}-\phi_3}{\psi_3}\right)^2 \\
+ & \quad \frac{1}{2}\left(\frac{\tilde{\beta}_l^{prop}-\phi_{2_l}}{\psi_{2_l}}\right)^2 + \frac{1}{2}\left(\frac{\tilde{\gamma}^{prop}-\phi_3}{\psi_3}\right)^2 \\
- & \quad \frac{1}{2}\left(\begin{bmatrix}\tilde{\beta}_l^{old}\\ \tilde{\gamma}^{old}\end{bmatrix}-\begin{bmatrix}\tilde{\beta}_l^{prop}\\ \tilde{\gamma}^{prop}\end{bmatrix}\right)^T \mathbf{\Sigma}_{[2_l,3]}^{-1}\left(\begin{bmatrix}\tilde{\beta}_l^{old}\\ \tilde{\gamma}^{old}\end{bmatrix}-\begin{bmatrix}\tilde{\beta}_l^{prop}\\ \tilde{\gamma}^{prop}\end{bmatrix}\right) \\
+ & \quad \frac{1}{2}\left(\begin{bmatrix}\tilde{\beta}_l^{prop}\\ \tilde{\gamma}^{prop}\end{bmatrix}-\begin{bmatrix}\tilde{\beta}_l^{old}\\ \tilde{\gamma}^{old}\end{bmatrix}\right)^T \mathbf{\Sigma}_{[2_l,3]}^{-1}\left(\begin{bmatrix}\tilde{\beta}_l^{old}\\ \tilde{\gamma}^{old}\end{bmatrix}-\begin{bmatrix}\tilde{\beta}_l^{prop}\\ \tilde{\gamma}^{prop}\end{bmatrix}\right).
\end{aligned}
$$

##### S 1.6.6 Hamiltonian Monte Carlo sampling scheme

The sampling method used here is the same as in the linear space, however we will define the Hamiltonian systems based upon the new system. Therefore, we now have $\tilde{\mathbf{q}} = \tilde{\theta}$, and the Hamiltonian function is

$$
H\left(\tilde{\mathbf{q}}, \mathbf{p}\right) = L\left(\tilde{\mathbf{q}}\right) + V\left(\tilde{\mathbf{q}}\right) + T\left(\mathbf{p}\right),
$$

with

$$
\begin{aligned}
L(\tilde{\mathbf{q}}) &= -\left(\log\left(\mathbb{P}(\tilde{\mathbf{q}})\right)\right) \\
&= \sum_{n=1}^{N} \log\left(\textbf{Normal}\left(\tilde{q}_n; \phi_n, \psi_n\right)\right), \\
&= \log\left(\psi\sqrt{2\pi}\right) + \frac{1}{2}\left(\frac{\tilde{q}_n - \phi_n}{\psi_n}\right)^2 \\
V(\tilde{\mathbf{q}}) &= -\sum_{k=1}^{K}\sum_{j=1}^{J_k} \log\left(\sum_{i=1}^{L} P^{t,\sigma_*^{old},\bar{b}^{old},10^{\tilde{\theta}^{prop}}}\left(\sigma_i, m_k^j\right)\right),
\end{aligned}
$$

and $T(\mathbf{P})$ is the same as in the linear space.

In order to advance a half step forward using $H^1(\tilde{\mathbf{q}}, \mathbf{p})$, we now use the change of variables $\tilde{\mathbf{q}} = \log_{10}(\mathbf{q})$ and find

$$
\frac{\partial V(\tilde{\mathbf{q}})}{\partial \tilde{\mathbf{q}}} = \frac{\partial V(\mathbf{q})}{\partial \mathbf{q}} \frac{\partial \mathbf{q}}{\partial \tilde{\mathbf{q}}} = 10^{\tilde{\mathbf{q}}} \log(10) \frac{\partial V(\mathbf{q})}{\partial \mathbf{q}},
$$

where $\frac{\partial V(\mathbf{q})}{\partial \mathbf{q}}$ is the same as $V_{\mathbf{q}}(\mathbf{q})$ above.

In order to advance the full step forward using $H^2(\tilde{\mathbf{q}})$ we use $L_{\tilde{\mathbf{q}}}(\tilde{\mathbf{q}})$, demonstrated for the rates $\tilde{\beta}_l$ and $\tilde{\gamma}$,

$$
L_{\tilde{\mathbf{q}}}(\tilde{\mathbf{q}}) = \begin{pmatrix} \frac{\tilde{\beta}_l - \phi_{2_l}}{\psi_{2_l}^2} \\ \frac{\tilde{\gamma} - \phi_3}{\psi_3^2} \end{pmatrix}.
$$

To approximate the system, we again apply a second order and symplectic implicit midpoint rule integration, using customMATLAB code, retrieved at [80]

$$
\frac{\tilde{\mathbf{q}}^{new} - \tilde{\mathbf{q}}^{old}}{h} = M^{-1}\left(\frac{\mathbf{p}^{old} + \mathbf{p}^{new}}{2}\right) \qquad \textbf{(3)}
$$

$$
\frac{\mathbf{p}^{new} - \mathbf{p}^{old}}{h} = L_{\tilde{\mathbf{q}}}\left(\frac{\tilde{\mathbf{q}}^{old} + \tilde{\mathbf{q}}^{new}}{2}\right).
$$

Solving for $\mathbf{P}^{new}$, and demonstrating with the parameter $\tilde{\beta}_l$, we find

$$
p_{2_l}^{new} = h\left(\frac{\left(\frac{\tilde{\beta}_l^{old} + \tilde{\beta}_l^{new}}{2}\right) - \phi_{2_l}}{\psi_{2_l}^2}\right) + p_{2_l}^{old}. \qquad \textbf{(4)}
$$

Plugging Eq. S (4) into Eq. S (3) we follow these algebraic steps,

$$
\begin{aligned}
\frac{\tilde{\beta}_l^{new} - \tilde{\beta}_l^{old}}{h} &= \frac{1}{m_{2_l}} \left( \frac{p_{2_l}^{old} + \left( h \frac{\left(\frac{\tilde{\beta}_l^{old}+\tilde{\beta}_l^{new}}{2}\right) - \phi_{2_l}}{\psi_{2_l}^2} + p_{2_l}^{old} \right)}{2} \right) \\
\frac{\tilde{\beta}_l^{new} - \tilde{\beta}_l^{old}}{h} &= \frac{1}{m_{2_l}} \left( p_{2_l}^{old} + \frac{h}{2\psi^2} \left( \frac{\tilde{\beta}_l^{old} + \tilde{\beta}_l^{new}}{2} - \phi_{2_l} \right) \right) \\
\tilde{\beta}_l^{new} - \tilde{\beta}_l^{old} &= \frac{h}{m_{2_l}} p_{2_l}^{old} + \frac{h^2}{2\psi^2 m_{2_l}} \left( \frac{\tilde{\beta}_l^{old} + \tilde{\beta}_l^{new}}{2} - \phi_{2_l} \right) \\
\tilde{\beta}_l^{new} - \tilde{\beta}_l^{old} &= \frac{h}{m_{2_l}} p_{2_l}^{old} + \frac{h^2}{4\psi^2 m_{2_l}} \tilde{\beta}_{2_l}^{old} + \frac{h^2}{4\psi^2 m_{2_l}} \tilde{\beta}_l^{new} - \frac{h^2}{2\psi^2 m_{2_l}} \phi_{2_l} \\
\tilde{\beta}_l^{new} - \frac{h^2}{4\psi^2 m_{2_l}} \tilde{\beta}_{2_l}^{new} &= \frac{h}{m_{2_l}} p_{2_l}^{old} + \frac{h^2}{4\psi^2 m_{2_l}} \tilde{\beta}_{2_l}^{old} - \frac{h^2}{2\psi^2 m_{2_l}} \phi_{2_l} + \tilde{\beta}_l^{old},
\end{aligned}
$$

to find

$$
\begin{aligned}
\tilde{\beta}_l^{new} &= \frac{\frac{h}{m_{2_l}} p_{2_l}^{old} - \frac{h^2}{2\psi_{2_l}^2 m_{2_l}} \phi_{2_l} + \left(1 + \frac{h^2}{4\psi_{2_l}^2 m_{2_l}}\right) \tilde{\beta}_l^{old}}{\left(1 - \frac{h^2}{4\psi_{2_l}^2 m_{2_l}}\right)} \\
&= \frac{\frac{4\psi_{2_l}^2 h}{m_{2_l}} p_{2_l}^{old} - \frac{2h^2}{m_{2_l}} \phi_{2_l} + \left(4\psi_{2_l}^2 + \frac{h^2}{m_{2_l}}\right) \tilde{\beta}_l^{old}}{\left(4\psi_{2_l}^2 - \frac{h^2}{m_{2_l}}\right)}.
\end{aligned}
$$

##### S 1.7 Robustness Analysis and Inference Validation on Synthetic Data

###### S 1.7.1 Maximum transcription rate

Due to the high number of model parameters, we analyze the method's sensitivity to changes in rates by varying the maximum production rate ($\beta_1$). If all other rates are maintained, changing $\beta_1$ corresponds to changing the expected number of RNA transcripts present in the cells. By varying $\beta_1$ from $\beta_1 = 0.3\ s^{-1}$ to $\beta_1 = 1\ s^{-1}$ (see Fig. S 1), we show that the method's prediction accuracy is robust under changes in the true gene network's associated parameters.

###### S 1.7.2 Specification of RNA degradation rate

In general, it is difficult to predict whether the amount of data provided to the method is sufficient to infer all rates simultaneously. To wit, Fig. S 2 shows how specifying one rate by hand allows smaller uncertainty in posteriors over the other rates.

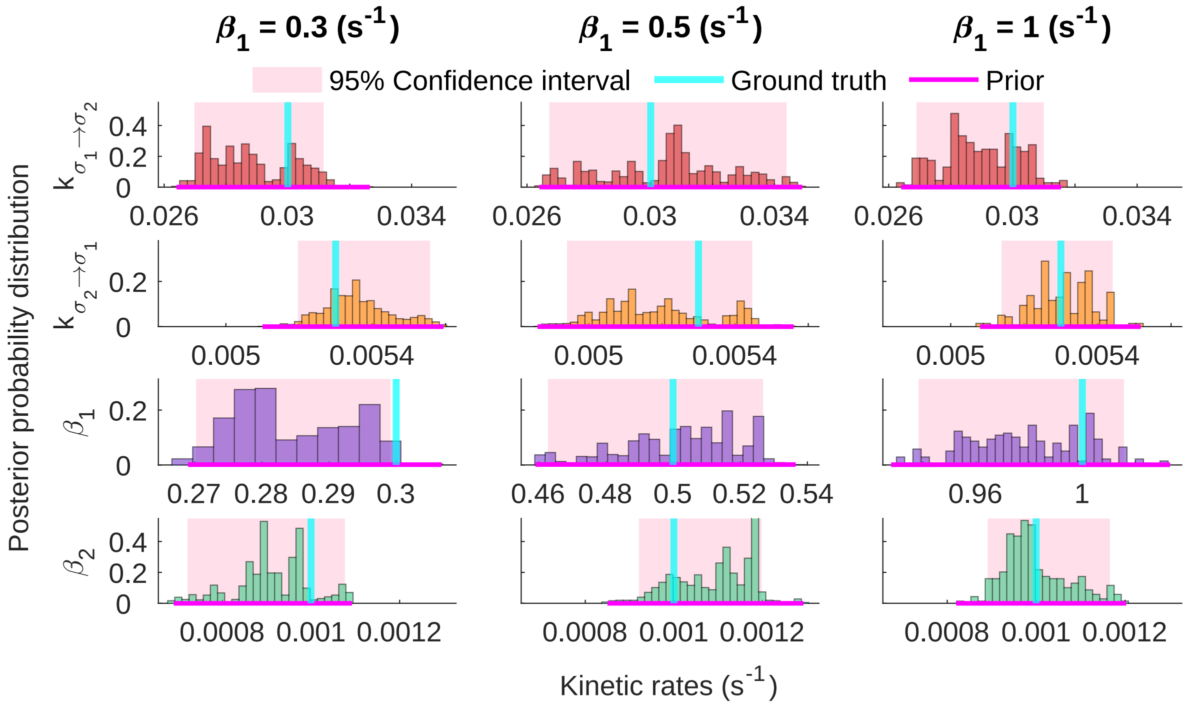


**Supplementary Figure 1.** ***Sensitivity analysis: transcription rate***. *Here, we show posterior distributions over: production rates* $\beta_l$*, and transition rates* $k_{\sigma_l \to \sigma_{l'}}$*. In each column, we show the comparable breadth of distributions for a network with two gene states, under various maximum ground truth production rates. We again find that our posterior maximum closely matches the ground truth. As before, each data point was generated using the Gillespie stochastic simulation algorithm [52, 53]. Rates in each column are inferred for 600 cells observed per time point with 20 collection times at* $[0, 10, 20, 30, 40, 50, 60, 70, 80, 90, 120, 180, 240, 360, 480, 600, 1200, 3600]$ $(s)$.

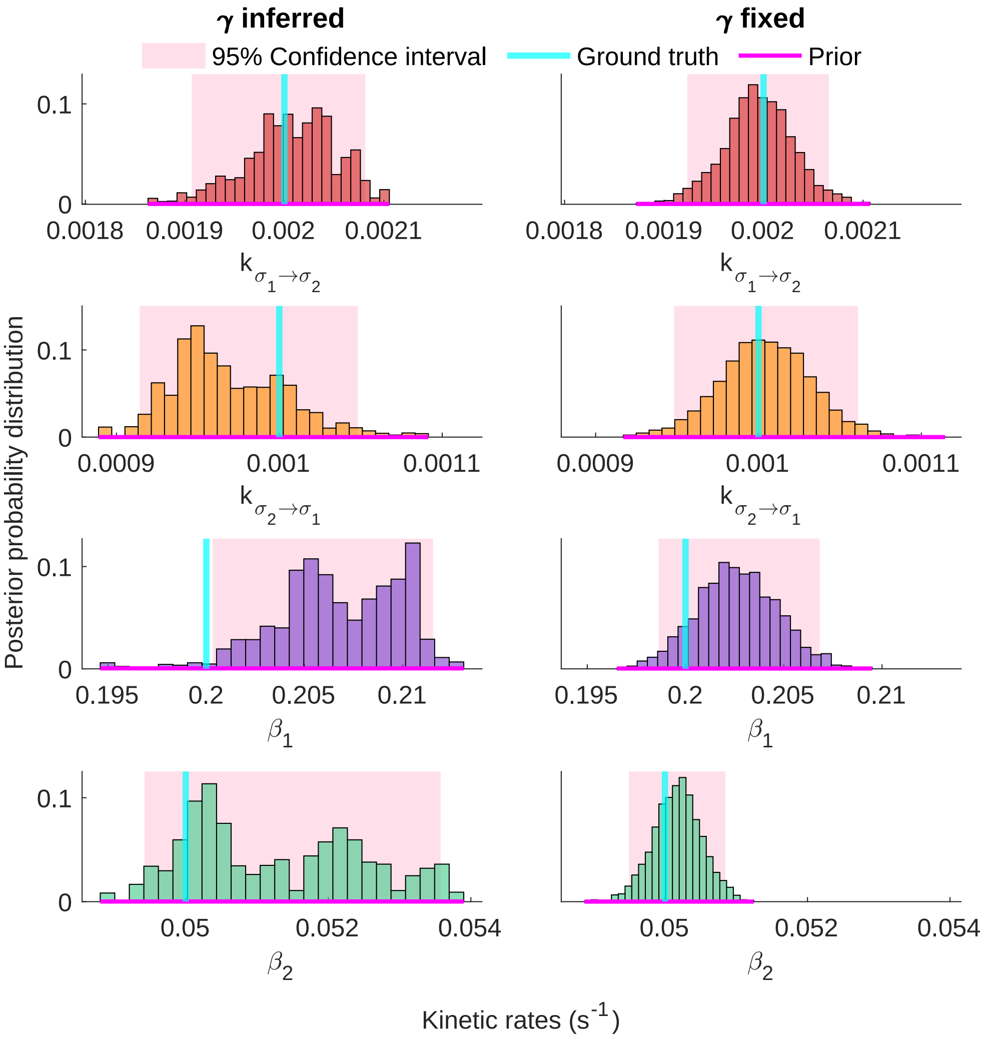


**Supplementary Figure 2.** ***Degradation rate: assumed vs inferred***. *Comparison of successful inference performed by either inferring, as shown in the left column, or fixing, as shown in the right column, the degradation rate ($\gamma$) for identical data from the three gene state model. As before, each row shows posteriors over a different rate, and the omitted rates are shown in Fig. S 13.*

### S 2 Supplementary Figures

#### S 2.1 Predictive Distributions and Supplemental Rate Histograms

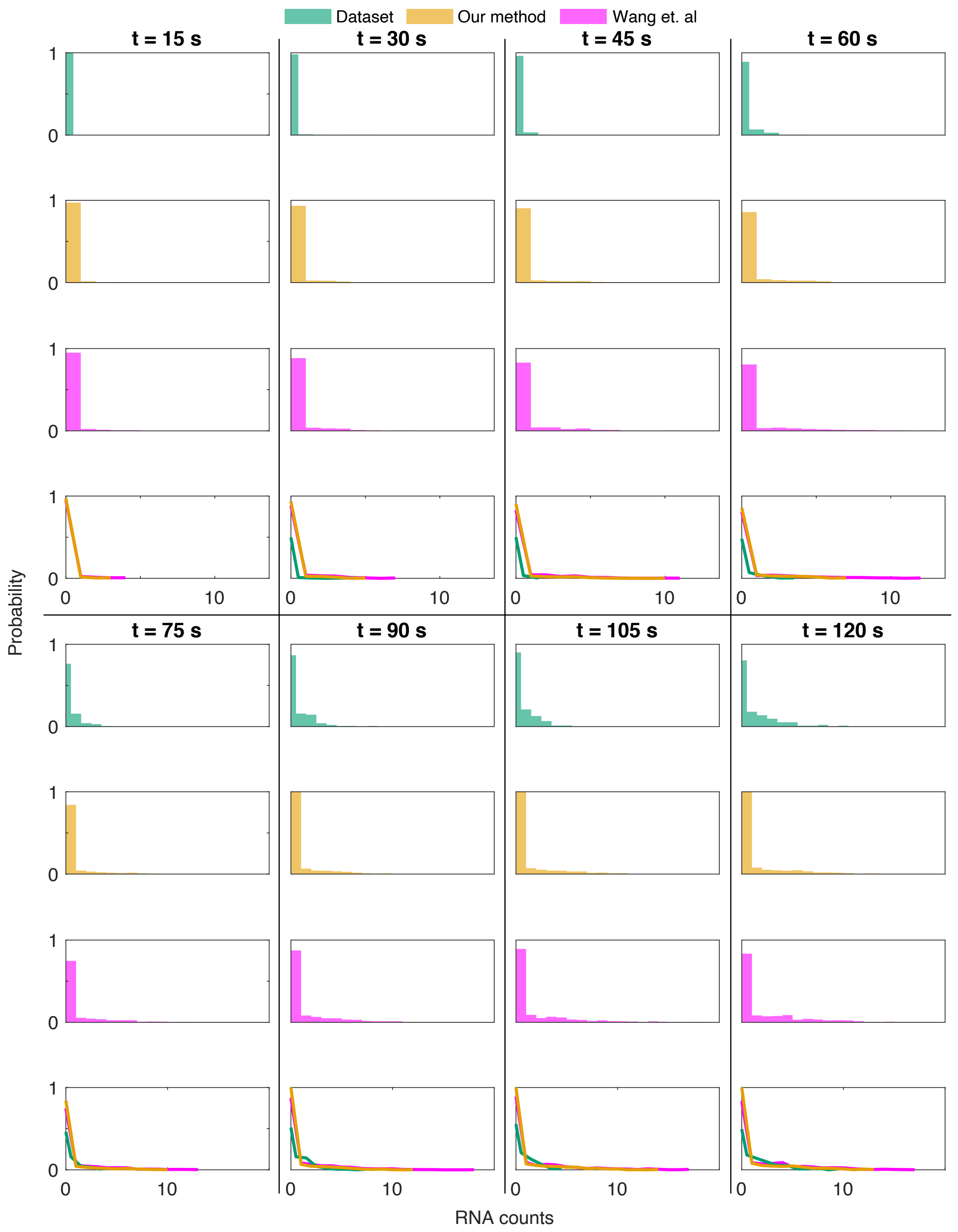


**Supplementary Figure 3. Predictive distributions corresponding to Fig. 4.** *Here we present the result of simulating 500 Gillespie trajectories per time point using "Our MAP" ($\theta'$ in Section 3.2) and results from [49] ("Wang et. al.") ($\theta$ in Section 3.2). The histograms above demonstrate that even when parameters differ widely, they make predict nearly identical RNA distributions. For this reason, it is important to avoid parameter estimates which only locally maximize their respective posterior.*

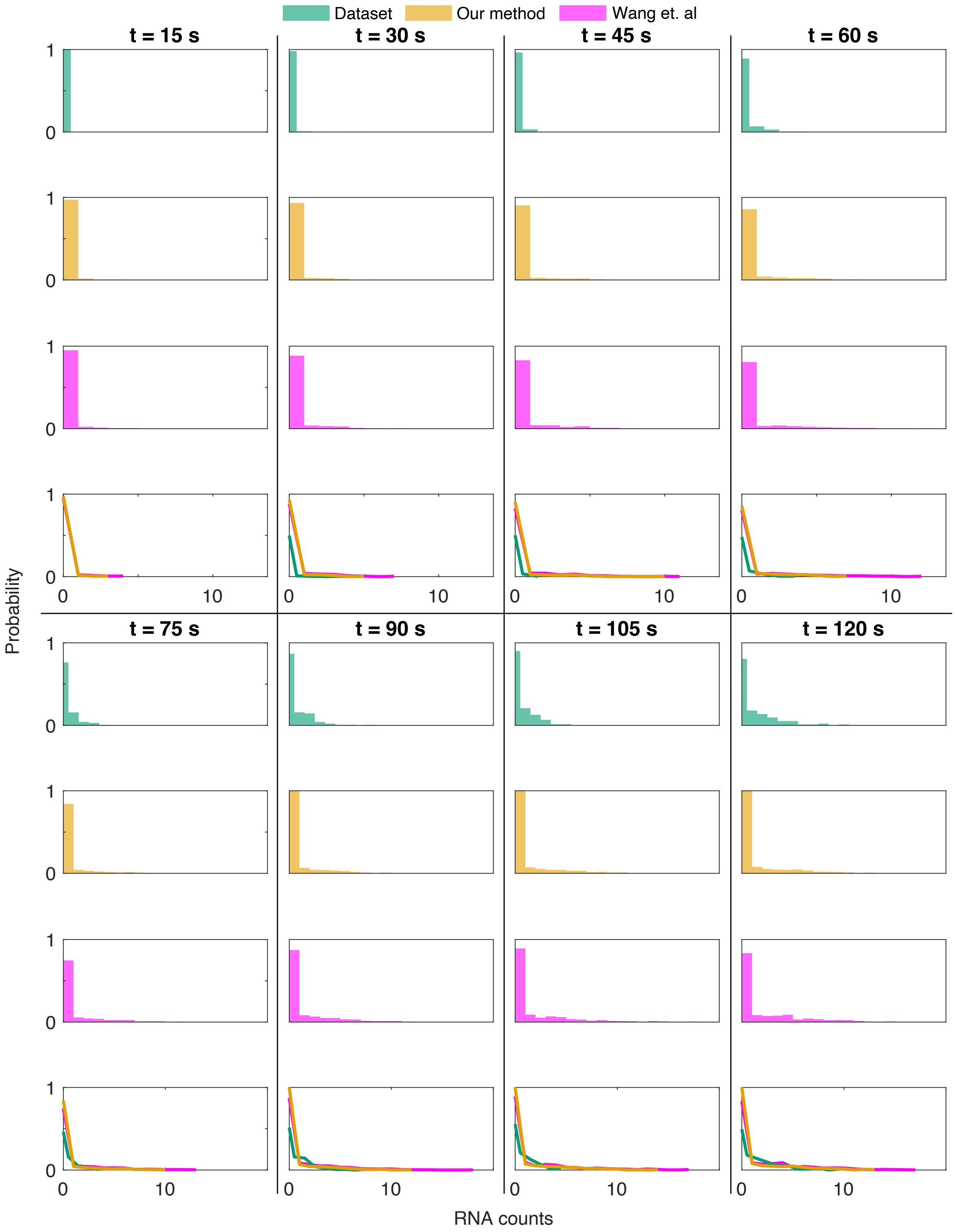


**Supplementary Figure 4. Predictive distributions corresponding to Fig. 4.** *Here we present the result of simulating 500 Gillespie trajectories per time point using "Our MAP" ($\theta'$ in Section 3.2) and results from [49] ("Wang et. al.") ($\theta$ in Section 3.2). See Fig. S 3 for context.*

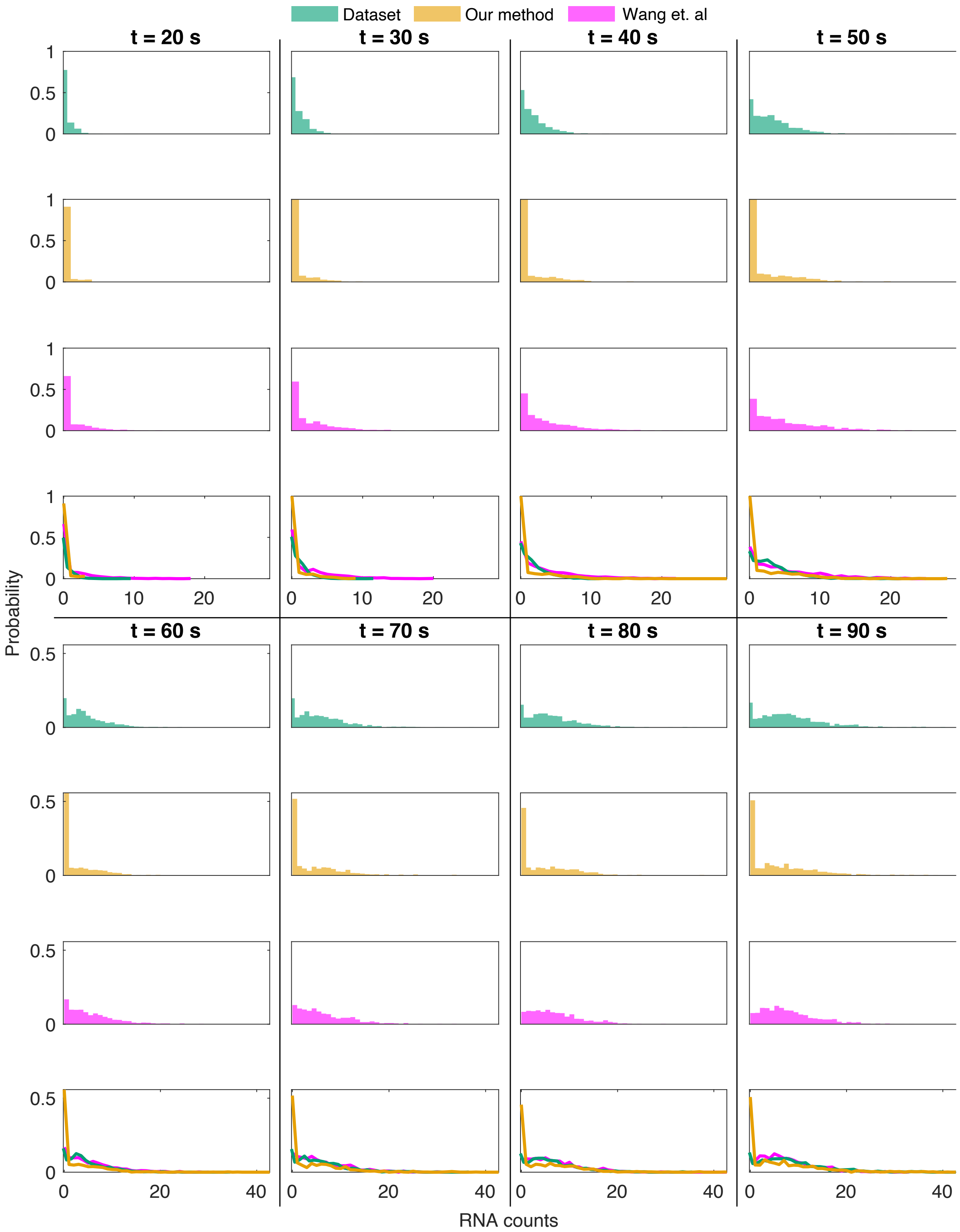


**Supplementary Figure 5. Predictive distributions corresponding to Fig. 5.** *Here we present the result of simulating 500 Gillespie trajectories per time point using "Our MAP" ($\theta'$ in Section 3.2) and results from [49] ("Wang et. al.") ($\theta$ in Section 3.2). See Fig. S 3 for context.*

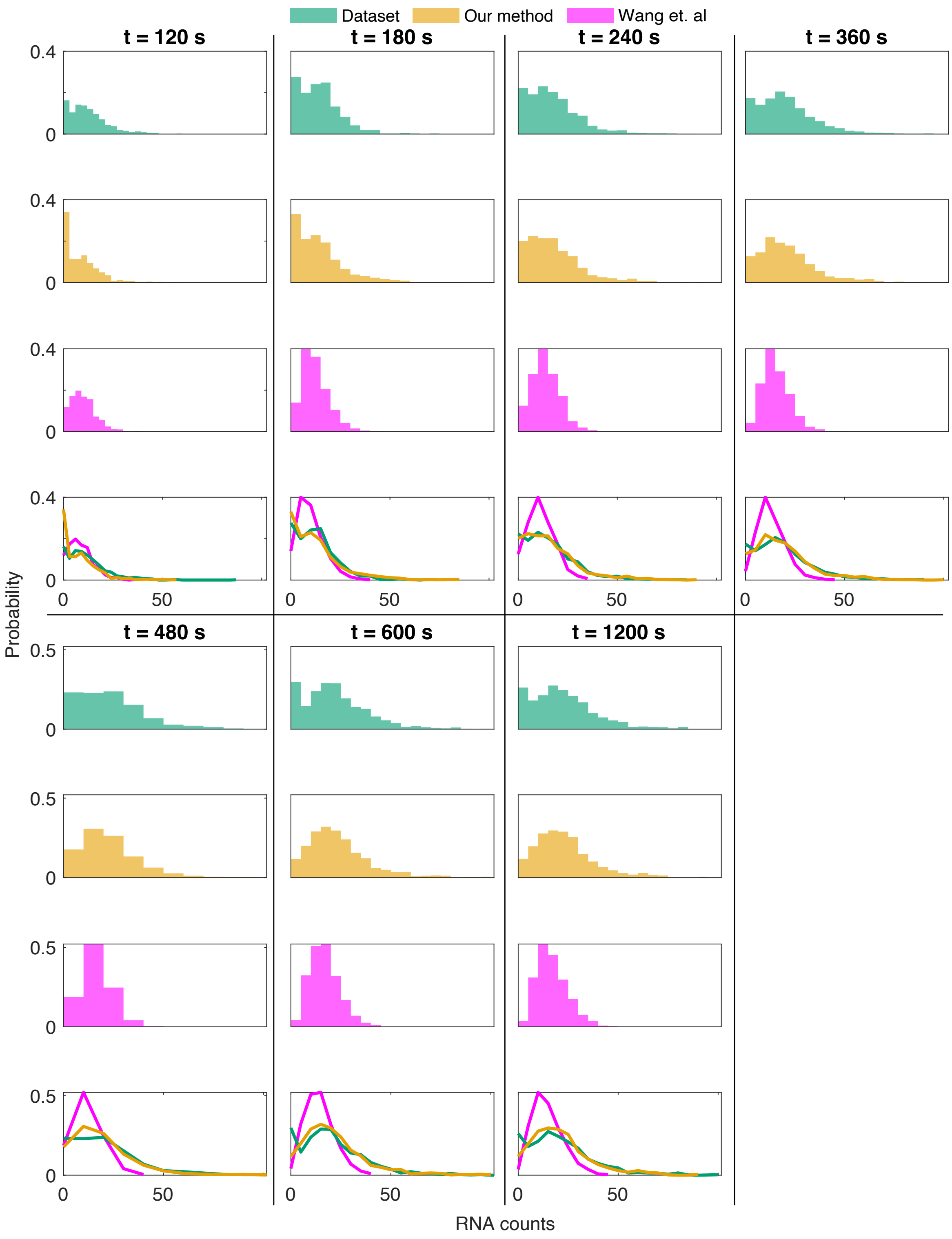


**Supplementary Figure 6. Predictive distributions corresponding to Fig. 5.** *Here we present the result of simulating 500 Gillespie trajectories per time point using "Our MAP" ($\theta'$ in Section 3.2) and results from [49] ("Wang et. al.") ($\theta$ in Section 3.2). See Fig. S 3 for context.*

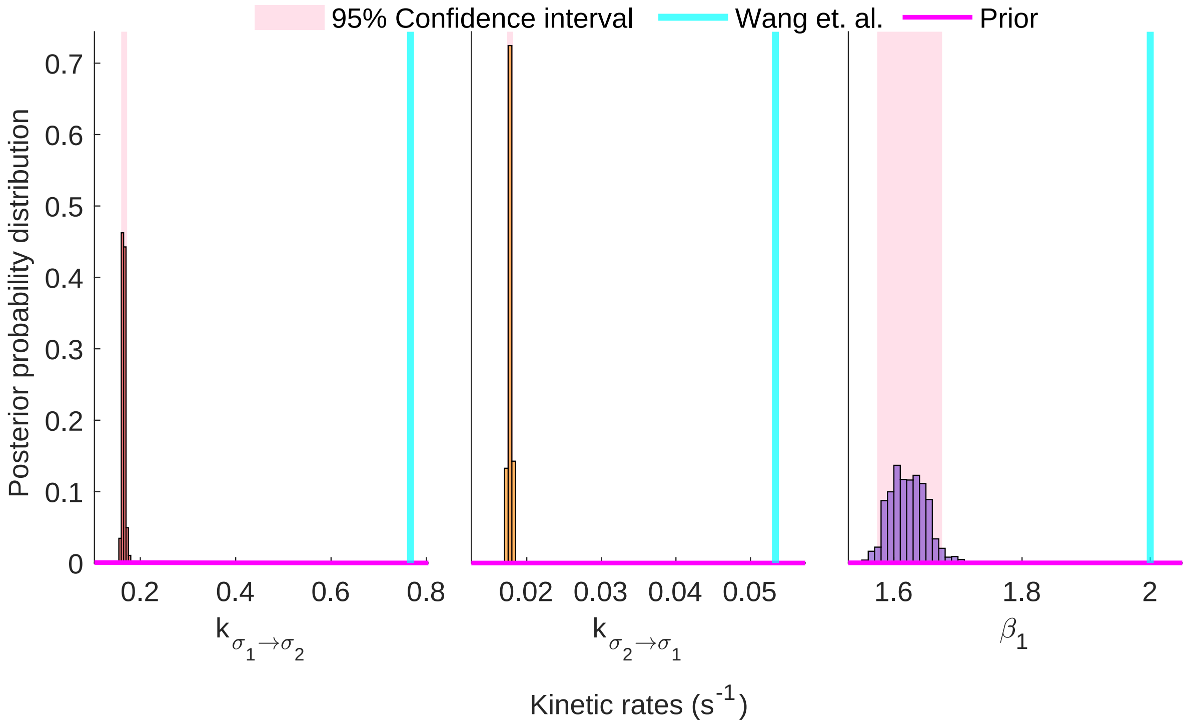


**Supplementary Figure 7.** ***Parametric inference on Escherichia coli data.*** *Here we show a comparison between the rates predicted when our model is constrained to only making predictions for the two gene state model, with one nontranscribing state. Owing to the overfitting problems associated with comparing likelihoods between models, and since we would still like to compare likelihoods, we obtained a parametric estimate within the model of [49]. The likelihoods of our MAP estimate ($\theta'$) and that of Wang et. al. [49] are compared in Section 3.2.1*

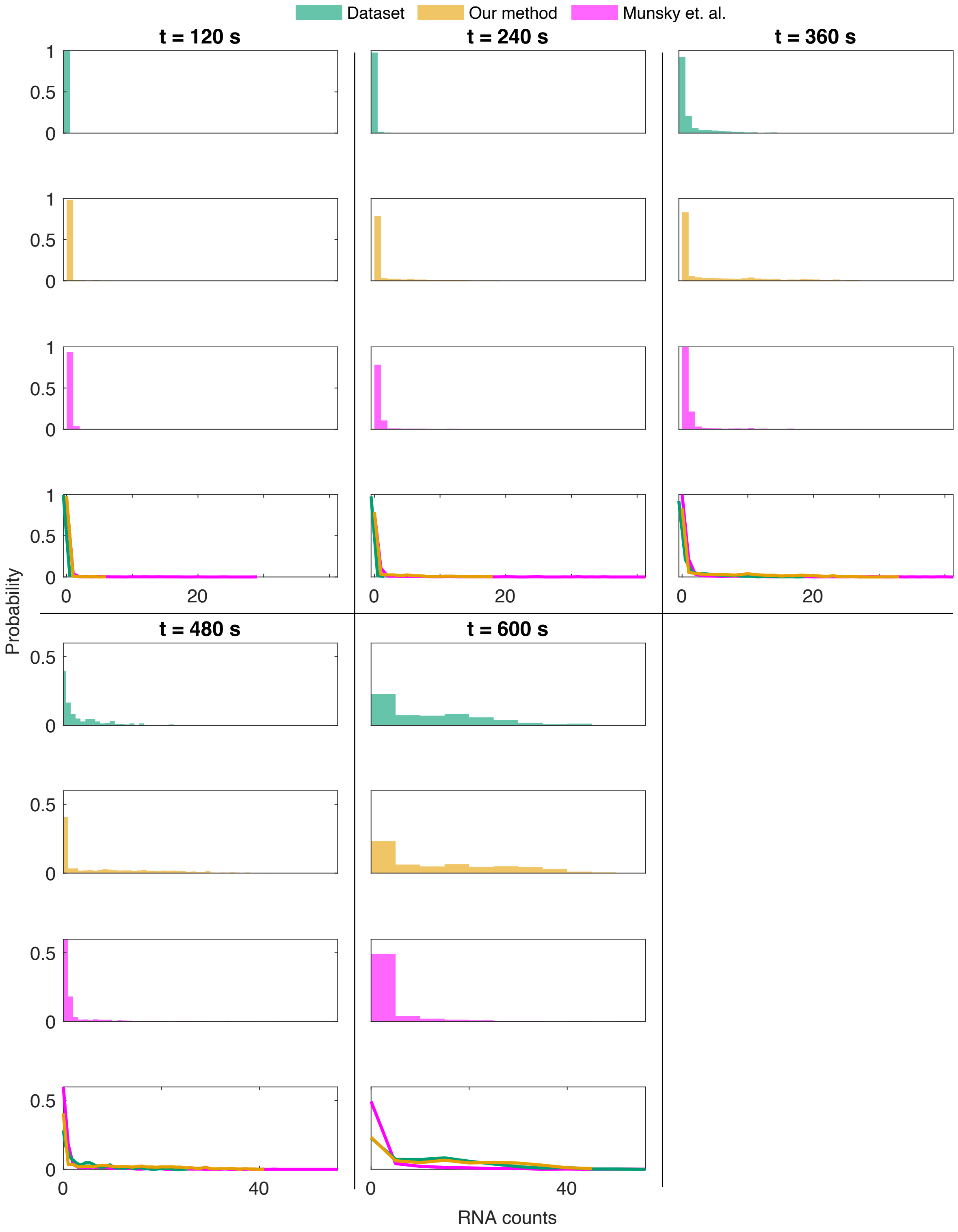


**Supplementary Figure 8. Predictive distributions corresponding to Fig. 6.** *Here we present the result of simulating 500 Gillespie trajectories per time point using "Our MAP" ($\theta'$ in Section 3.2) and resultd from [39] ("Munsky et. al.") ($\theta$ in Section 3.2). See Fig. S 3 for context.*

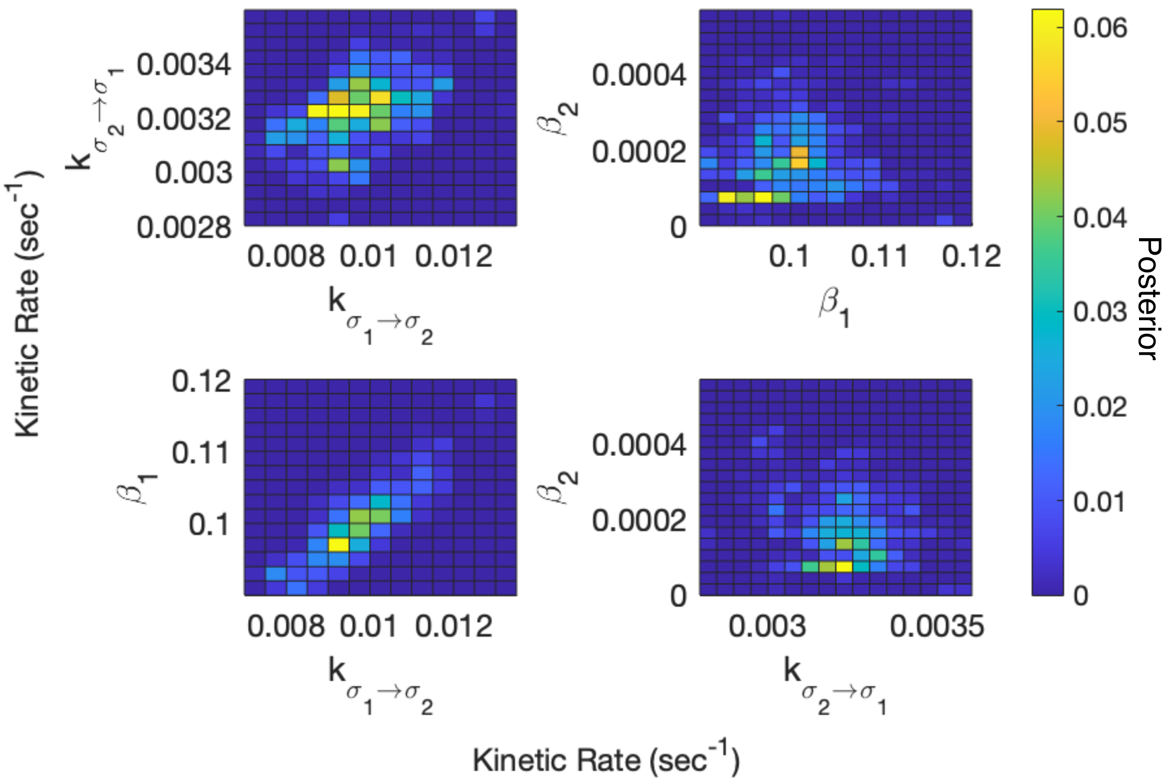


**Supplementary Figure 9. Joint histograms of rates for the three state model of Fig. 4.** *In this and subsequent joint histograms, we see examples of the type of local maxima described in Section 3.2.1 and Section 3.2.2. Though there may be significant density associated with regions away differ from our MAP estimate, these peaks do not globally maximize the posterior.*

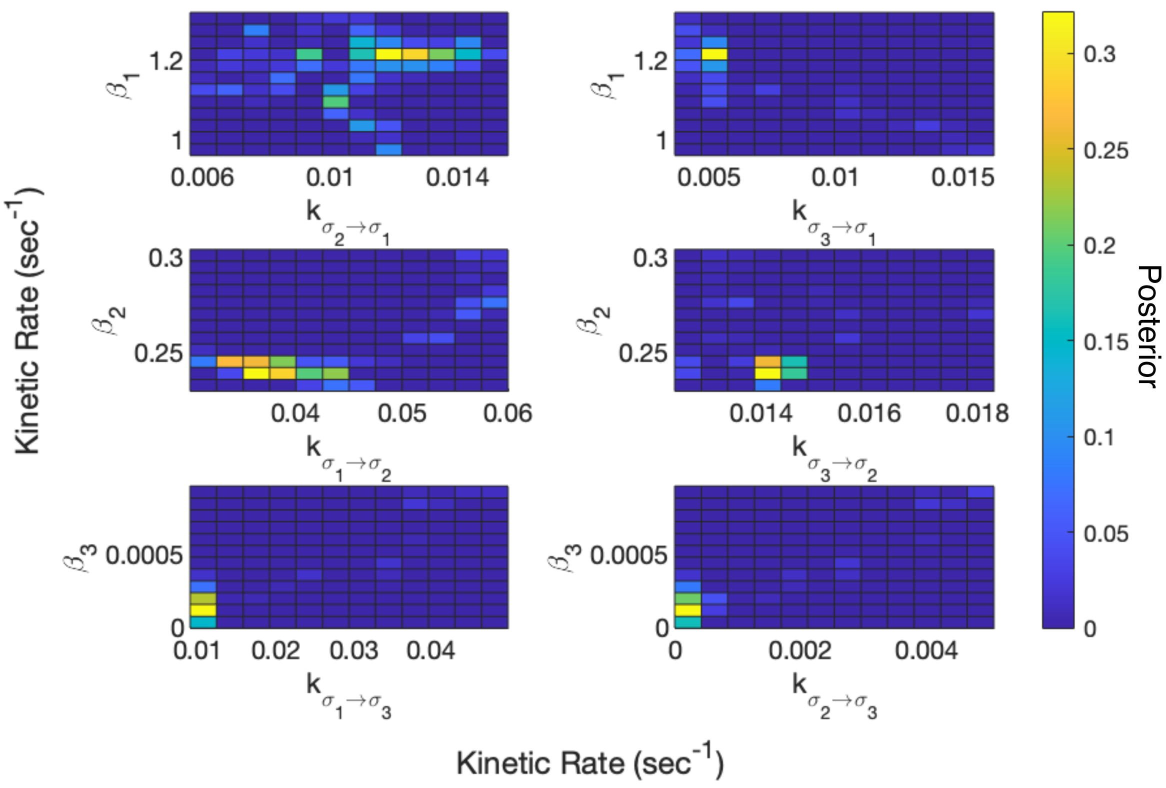


**Supplementary Figure 10.** **Joint histograms of rates for the three state model of Fig. 5.** *See Fig. S 9 for context.*

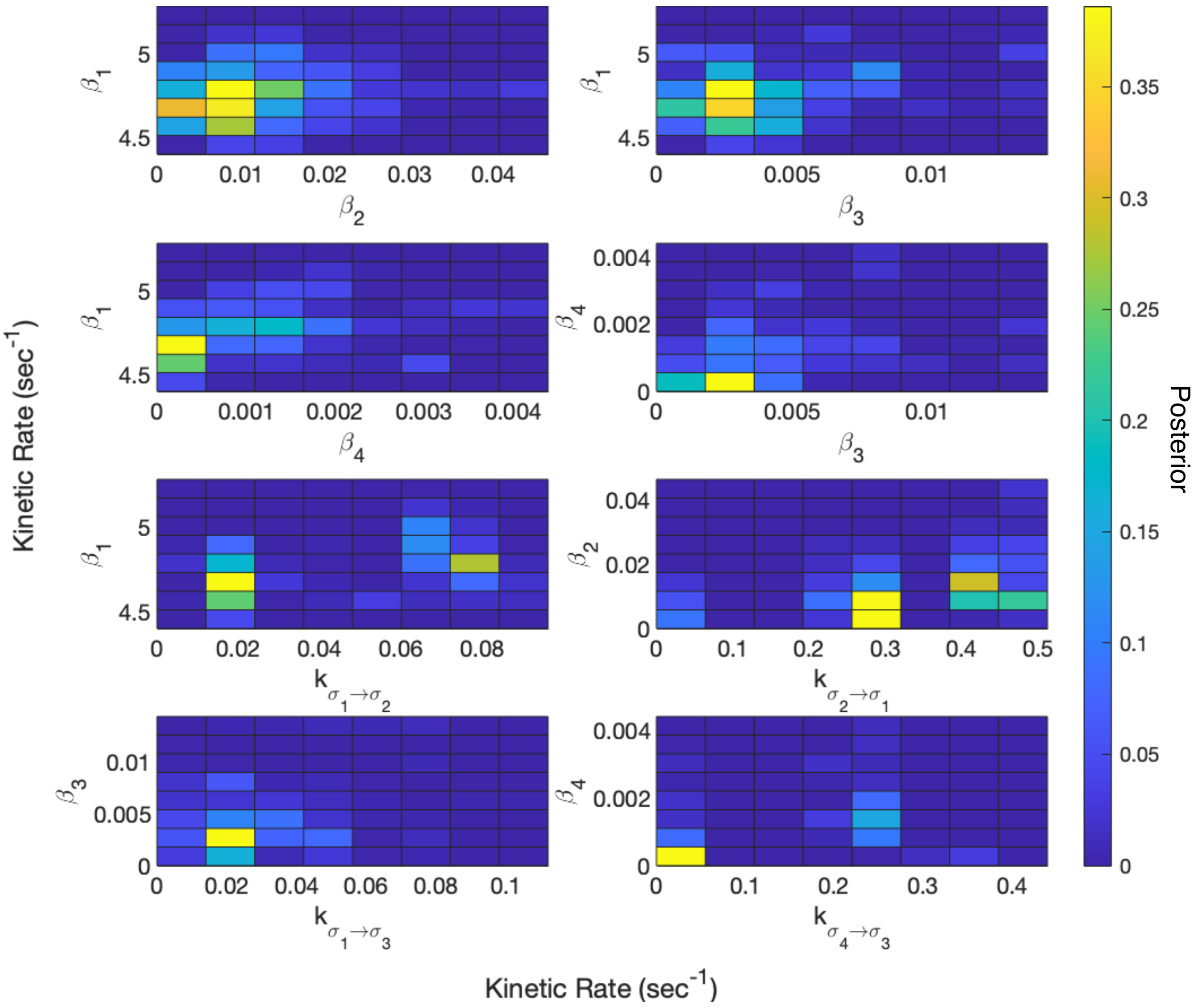


**Supplementary Figure 11.** **Joint histograms of rates for the four state model of Fig. 6.** *See Fig. S 9 for context.*

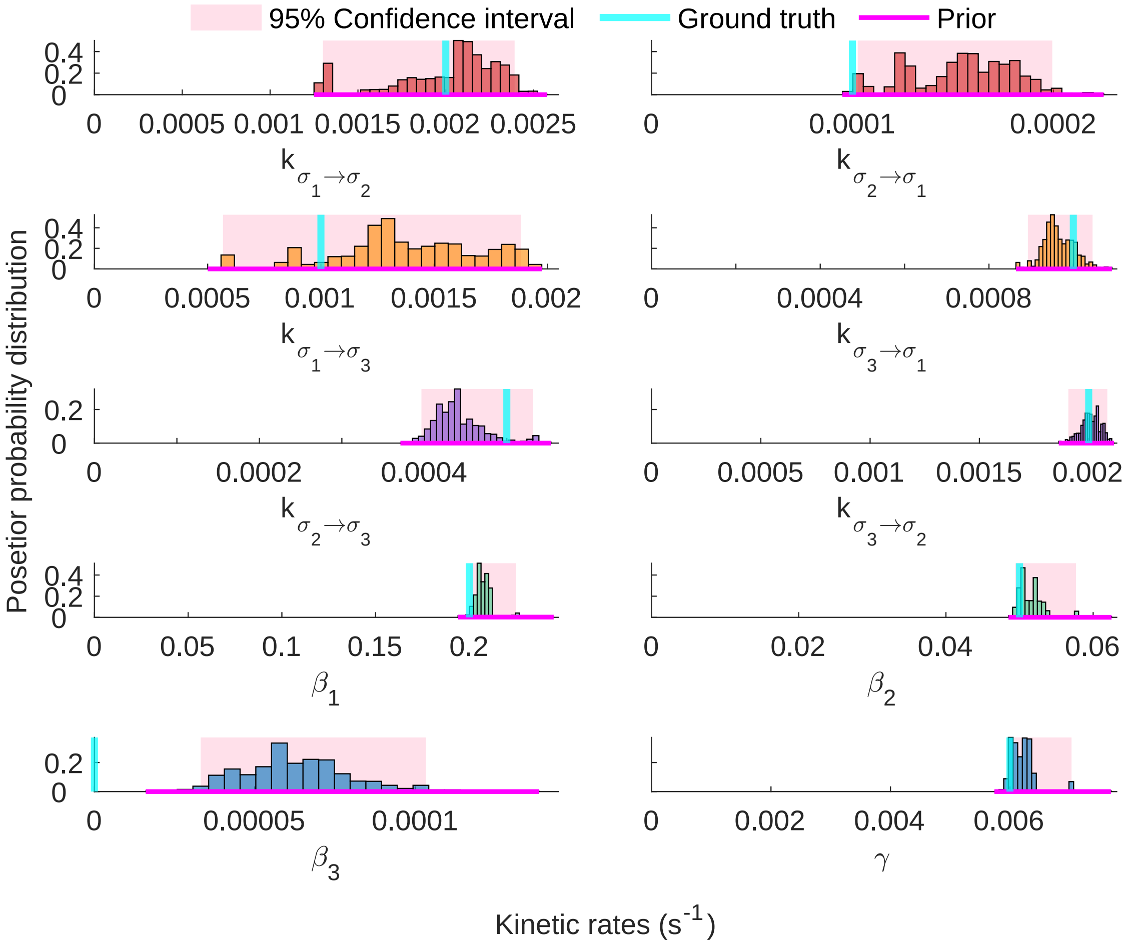


**Supplementary Figure 12. Histograms of additional rates for the three state model of Fig. 2.**

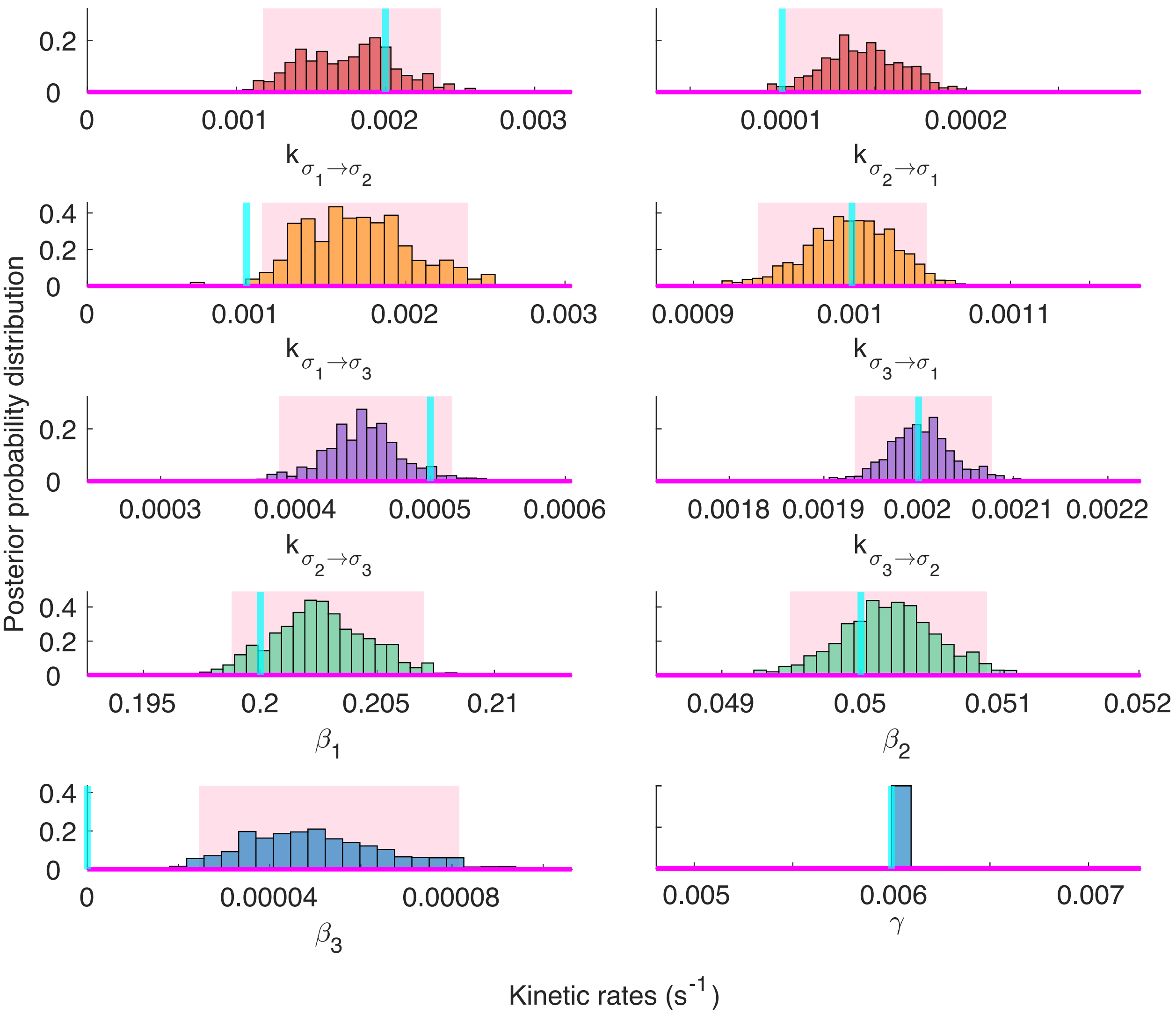


**Supplementary Figure 13.** **Histograms of additional rates for $\gamma$ fixed as in Fig. S 2.**
